# Genome sequences of the five *Sitopsis* species of *Aegilops* and the origin of polyploid wheat B-subgenome

**DOI:** 10.1101/2021.07.05.444401

**Authors:** Lin-Feng Li, Zhi-Bin Zhang, Zhen-Hui Wang, Ning Li, Yan Sha, Xin-Feng Wang, Ning Ding, Yang Li, Jing Zhao, Ying Wu, Lei Gong, Fabrizio Mafessoni, Avraham A. Levy, Bao Liu

**Author notes:** **Correspondence**: Lin-Feng Li, Avraham A. Levy, Bao Liu. These authors contributed equally to this work (Lin-Feng Li, Zhi-Bin Zhang).

## Abstract

Bread wheat (*Triticum aestivum* L., BBAADD) is a major staple food crop worldwide. The diploid progenitors of the A- and D-subgenomes have been unequivocally identified, that of B however remains ambiguous and controversial but is suspected to be related to species of *Aegilops*, section *Sitopsis*. Here, we report the assembly of chromosome-level genome sequences of all five *Sitopsis* species, namely *Ae. bicornis, Ae. longissima, Ae. searsii, Ae. sharonensis*, and *Ae. speltoides*, as well as partial assembly of *Ae. mutica* genome for phylogenetic analysis. Our results support that the donor of bread wheat B-subgenome is a distinct, probably extinct, diploid species that diverged from an ancestral progenitor of the B-lineage similar to *Ae. mutica* and *Ae. speltoides*. The five *Sitopsis* species have variable genome sizes (4.11-5.89 Gb) with high proportions of repetitive sequences (85.99-89.81%); nonetheless, they retain high collinearity with other wheat genomes. Differences in genome size are primarily due to independent post-speciation amplification of transposons rather than to inter-specific genetic introgression. We also identified a set of *Sitopsis* genes pertinent to important agronomic traits that can be harnessed for wheat breeding. These resources provide a new roadmap for evolutionary and genetic studies of the wheat group.

**Significance:** The origin of the B-subgenome of hexaploid bread wheat remains unknown. Here we report the assembly of chromosome-level genome sequences of all five *Sitopsis* species of the genus *Aegilops*, which are previously considered as possible direct progenitors or contributors to the B-subgenome. Our comparative genomic analyses reveal that the B-subgenome originated from an unknown, most likely extinct species phylogenetically distinct from *Ae. speltoides*, its extant closest relative. We also provide evidence that *Ae. speltoides* is neither the direct progenitor of the G-subgenome of tetraploid wheat *Triticum timopheevii*. The high-quality *Sitopsis* genomes provide novel avenues to identify new important genes for wheat breeding.

## Introduction

Hexaploid bread wheat (*Triticum aestivum* L., 2n = 6*x* = 42, BBAADD) is the most widely grown and largest acreage crop in the world, providing about 20% of the global calories and proteins in human consumption^1^. Bread wheat contains three closely related subgenomes (A, B and D) donated by distinct diploid species, which were reunited via a recent allohexaploid speciation event between a cultivated tetraploid wheat (genome BBAA) and a diploid goat grass (*Ae. tauschii*, DD) less than 10,000 years ago^2,3^. The cultivated tetraploid wheat, *T. turgidum* ssp. *durum* or ssp. *dicoccon* was domesticated from wild emmer wheat (*T. turgidum* ssp. *dicoccoides*, BBAA). Wild emmer wheat itself was formed via an earlier allotetraploidization event (<0.8 million years ago, MYA) between two wild diploid species of *Triticum/Aegilops*, which donated the A- and B-subgenomes, respectively^4,5^. It is established that the A- and D-subgenomes of polyploid wheat are derived from wild diploid wheat *T. urartu* (AA) and goat-grass *Ae. tauschii* (DD), respectively^6,7^. However, the origin of polyploid wheat B-subgenome remains unclear and controversial.

The hypothesis that the polyploid wheat B-subgenome originated from a diploid *Aegilops* species of section *Sitopsis* has been proposed since the 1950s. This inference was mainly based on the close morphological similarity of spikelet and karyotype structure between the *Sitopsis* species (*i.e*., *Ae. speltoides*) and species of the *Triticum* genus^8,9^. Yet, the comparisons of chromosome structure and meiotic pairing behavior revealed virtually absence of homologous synapsis between the polyploid wheat B-subgenome and all *Sitopsis* species genomes^10–12^. Molecular phylogeny and population genetic inferences showed either high genetic similarity^13^ or the closest phylogenetic relationship^5,14–16^ of *Ae. speltoides* to the wheat B-subgenome. These molecular and cytological evidences have led to the monophyletic origin hypothesis purporting that the wheat B-subgenome evolved from *Ae. speltoides* or a closely related species, but was modified at the polyploid level^8,10,12,17^. An alternative hypothesis is that the origin of the modern wheat B-subgenome or its diploid progenitor is polyphyletic, being shaped by hybridization or introgression of diverse genomic sequences from different *Triticum/Aegilops* species^18–20^. This scenario appeared to be congruent with transcriptome-based phylogenetic inferences that revealed frequent interspecific hybridizations in the species complex^21^. However, genome-scale evidence supporting the monophyletic or polyphyletic origins of the polyploid wheat B-subgenome is lacking.

The reference genomes of both hexaploid bread wheat and tetraploid wild emmer/cultivated durum wheats, as well as their diploid progenitors, *Ae. tauschii* and *T. urartu*, have been released in recent years^22–27^, but whole genome sequence of the diploid species related to the B-subgenome of polyploid wheat is not yet available. Here, we report chromosome-level genome assemblies of all the five diploid species of *Aegilops*, section *Sitopsis, i.e., Ae. bicornis, Ae. longissima, Ae. searsii, Ae. sharonensis*, and *Ae. speltoides*, as well as partial assembly of *Ae. mutica* genome. The reference-quality genome assemblies of these diploid species, together with those available for polyploid wheat and the A- and D-subgenome diploid progenitor species, provide a comprehensive repertory of genome resources for deeper evolutionary studies in the *Triticum*/*Aegilops* species complex. Our results also shed new light on the evolution of the B-lineage, confirming that *Ae. mutica* is the present-day species most close to the ancestral progenitor of the B-lineage^21^ and supporting that the donor of the polyploid wheat B-subgenome was a single diploid species, probably extinct, that is most closely related to, but also distinct from, the still extant *Ae. speltoides*. In addition, we show that the novel genomic resources from the *Sitopsis* can be mined to identify new genes and natural allele variants that can be utilized to cope with the ever-increasing global demand for wheat improvement.

## Results

### Sequence assemblies and genome features

Identities of the five diploid *Sitopsis* species (2n = 2*x* = 14) of *Aegilops* were confirmed by fluorescence *in situ* hybridization (FISH) and spike morphology (**Supplementary Fig. 1**). The same, multi-generation-selfed, individual (bagged) of each species was used for genome sequencing and assembly. Sizes of the assembled genomes of the five species ranged from 4.11 to 5.89 Gb, broadly consistent with values (4.45-6.02 Gb) estimated by flow cytometry (**Table 1**). Notably, *Ae. speltoides* (4.11 Gb) has the smallest genome among the five species, which is close in size to those of the bread wheat D-subgenome (3.95 Gb)^25^ and its donor *Ae. tauschii* (4.30 Gb)^24^. In contrast, the remaining four *Sitopsis* species, *Ae. bicornis* (5.64 Gb), *Ae. longissima* (5.80 Gb), *Ae. searsii* (5.34 Gb) and *Ae. sharonensis* (5.89 Gb), all have much larger genomes similar to the polyploid wheat A- (4.86-4.94 Gb) and B-subgenomes (5.11-5.18 Gb)^22,25,26^, and to the genome of *T. urartu* (4.94 Gb)^23^.

**Table 1.** Statistics of genome features of the five *Sitopsis* species.

|  | Assembly parameters | *Ae. bicornis* | *Ae. longissima* | *Ae. searsii* | *Ae. sharonensis* | *Ae. speltoides* |
| --- | --- | --- | --- | --- | --- | --- |
| Genome assembly | Genome size* (Gb) | 5.54 | 6.02 | 5.37 | 5.95 | 4.45 |
|  | Total length of contigs (Gb) | 5.64 | 5.80 | 5.34 | 5.89 | 4.11 |
|  | GC content (%) | 46.39 | 46.40 | 46.01 | 46.24 | 46.34 |
|  | N50 length (contig) (Mb) | 8.72 | 1.05 | 0.56 | 1.01 | 1.78 |
| Repetitive sequence | Retrotransposons (Gb) | 3.99 | 4.15 | 3.55 | 4.21 | 2.94 |
|  | DNA transposons (Gb) | 1.01 | 0.89 | 1.03 | 0.91 | 0.52 |
|  | Total (Gb) | 5.08 | 5.11 | 4.64 | 5.19 | 3.54 |
| Protein-coding genes | Predicted protein-coding gene | 61,354 | 63,326 | 62,804 | 61,849 | 61,084 |
|  | High confidence gene | 40,222 | 37,201 | 37,995 | 38,440 | 37,607 |
|  | Average transcript length (bp) | 1,484 | 1,657 | 1,554 | 1,627 | 1,622 |
|  | Average CDS length (bp) | 1,193 | 1,293 | 1,290 | 1,289 | 1,319 |
|  | Average exon length (bp) | 325 | 358 | 329 | 351 | 348 |
|  | Average intron length (bp) | 507 | 527 | 860 | 818 | 834 |
|  | Functionally annotated | 37,082 | 36,539 | 37,489 | 37,798 | 37,301 |
|  | BUSCO integrity (%) | 97.99 | 94.03 | 92.92 | 93.19 | 93.75 |

Note: \*, Genome size was estimated by flow cytometry.

To determine the origin of the variable genome content, we annotated both protein-coding genes and repetitive sequences of the five species and compared with the other relevant wheat species, including *Ae. tauschii* and *T. urartu*, wild emmer, domesticated durum and bread wheat. A total of 37,201-40,222 high confidence protein-coding genes were predicted in the five *Sitopsis* genome assemblies, with >92.2% of which being functionally annotated in the GO/KEGG/KOG/NR databases (**Table 1**). On average, the genes of the *Sitopsis* species encode transcripts of 1,193-1,319 bp in length, which are comparable to the three bread wheat subgenomes (1,310-1,351 bp) and *Ae. tauschii* (1,144 bp) but longer than to *T. urartu* (998 bp)^23–25^. In addition, our results show that difference in genome size among the five *Sitopsis* species is mainly attributed to the variable total length of repetitive sequences (3.54-5.19 Gb, 86.13-88.11% of the total), including 2.94-4.21 Gb (66.48-71.47%) retrotransposons and 0.52-1.03 Gb (12.54-19.21%) DNA transposons (**Table 1**).

Distribution patterns of GC content, protein-coding genes and repetitive sequences were assessed for the five *Sitopsis* species and bread wheat B-subgenome. Broadly consistent with previously published genomes of wheat species, all five *Sitopsis* species show higher gene density and lower GC content in distal than proximal chromosomal regions (**Fig. 1a-c**). A general genomic feature of the repetitive sequences of the five species and bread wheat B-subgenome is that *copia-like* long-terminal repeat (LTR) retrotransposons tend to cluster at telomeric regions of all seven chromosomes (**Fig. 1d**), whereas a reverse distribution pattern was observed in the *gypsy-like* LTR retrotransposons (**Fig. 1e**). It is notable that *Ae. speltoides* shows distinct distribution density of *copia-like* retrotransposons and *CACTA* DNA transposons compared to bread wheat B-subgenome and the other four *Sitopsis* species across all seven chromosomes (**Fig. 1d** and **f**). Nevertheless, estimates of the overall unique *k*-mer frequency revealed similar density of repetitive sequence between the five *Sitopsis* species and wheat B-subgenome (**Fig. 1g**), but far lesser than the A- and D-subgenomes detailed in previous study^28^ (Kruskal-Wallis test, *p* < 0.001). We also performed genome collinearity analyses to assess the differences in genome structure. Although these *Triticum*/*Aegilops* species contain large proportions of repetitive sequences and differ substantially in genome sizes, they still retain high collinear genomes (**Supplementary Fig. 2**). Notably, two previously identified species-specific translocation events, 4A/5A/7B in tetraploid/hexaploid wheat^29^ and 7S^l^/4S^l^ translocation in *Ae. longissima*^12,30^, were confirmed in our genome collinear analyses, corroborating the quality of our genome assemblies.

**Figure 1.**
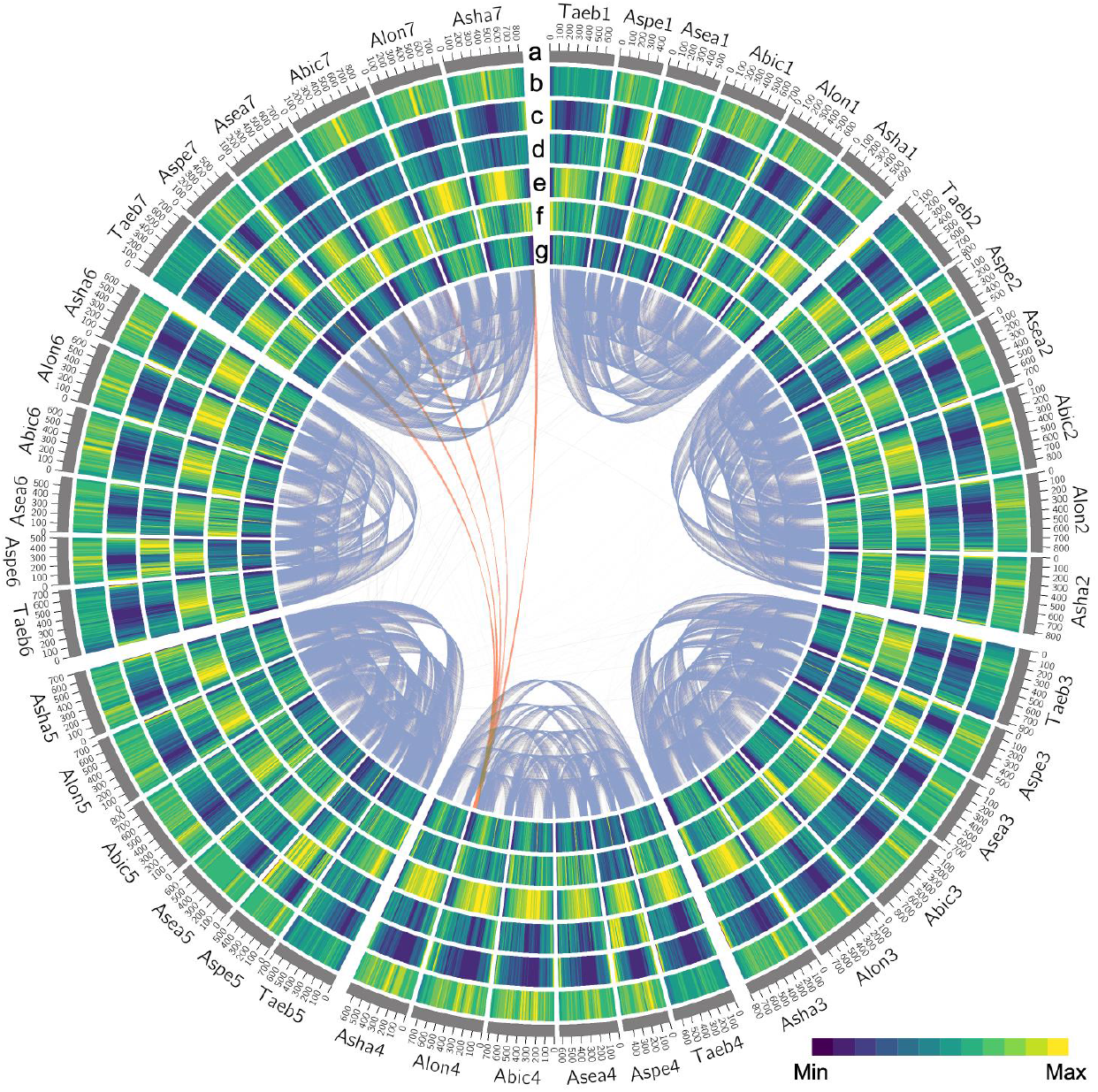
Structural, functional, and syntenic landscape of the five *Sitopsis* species and bread wheat B-subgenome. Circular diagram from outside to inside are: **a**, Species and chromosome names, each tick mar is 100 Mb in length. **b**, Percent of GC content. **c**, Density of high confidence gene (0-21 per Mb). **d**, *The copia-like* retrotransposon density. **e**, *The gypsy-like* retrotransposon density. **f**, *CACTA* DNA transposon density. **g**, Distribution of unique 20-mer frequencies across physical chromosomes (5-160 K-mer/Mb). Color of links is blue between homeologous chromosomes and orange in cases of large translocations. Note: Abic: *Ae. bicornis*, Alon: *Ae. longissima*, Asea: *Ae. searsii*, Asha: *Ae. shanronensis*, Aspe: *Ae. speltoides*, Taeb: B-subgenome of the bread wheat.

### Molecular phylogeny, divergence time and genetic similarity

Phylogenetic relationships of the five *Sitopsis* and other *Triticum*/*Aegilops* species were reconstructed based on single copy orthologous gene (SCOG), reduced representative genomic region (RRGR) and whole-genome single nucleotide polymorphism (SNP) datasets. Similar to previously inferred phylogenies^4,21^, the diploid species and polyploid wheat subgenomes fall into three independent clades corresponding to the A-, B- and D-lineages, with *Ae. speltoides* being clustered with polyploid wheat B-subgenome (B-lineage) while the rest four *Sitopsis* species being grouped with bread wheat D-subgenome (D-lineage) and its diploid donor *Ae. tauschii* (D-lineage) (**Fig. 2a** and **Supplementary Fig. 3**).

**Figure 2.**
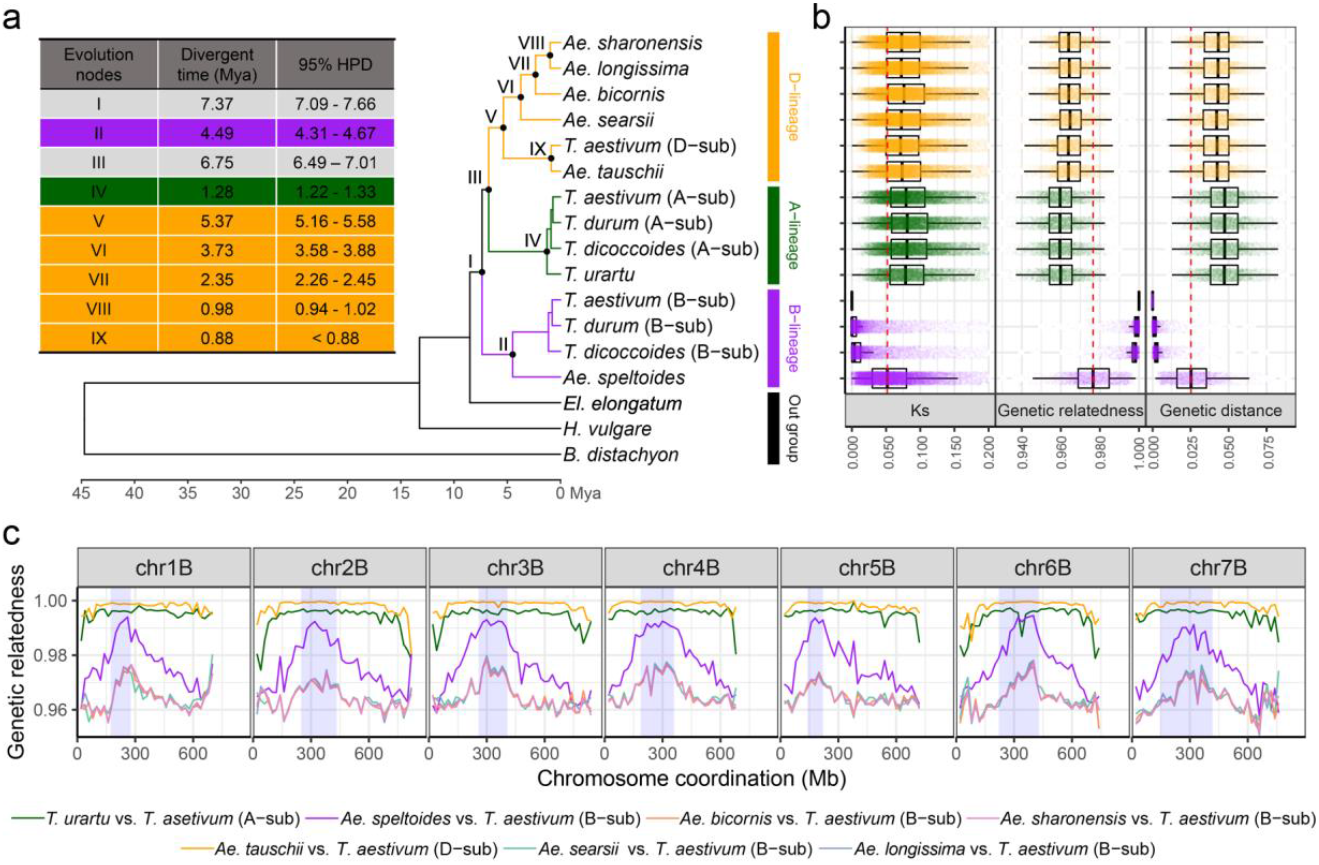
Phylogenetic relationships, divergence times and genetic similarities of the *Triticum*/*Aegilops* species complex. **a**, Maximum likelihood tree and divergence times of the *Triticum*/*Aegilops* species complex based on single copy orthologous genes. The green, purple and orange branches represent the A-, B- and D-lineages, respectively. Black color branches are the common ancestor of *Triticum*/*Aegilops* species and outgroup species. **b**, Synonymous mutation rate (Ks), genetic relatedness and genetic distance between the bread wheat B-subgenome and the other diploid species and polyploid wheat subgenomes based on collinear genes and reduced representative genomic regions. Green, purple and orange colors indicate these comparisons were obtained from the A-, B- and D-lineages. Numbers on the X-axis are the values of genetic similarity. Red dashed line indicates the mean values between *Ae. speltoides* and bread wheat B-subgenome. **c**, Distribution of the genetic relatedness between the five *Sitopsis* species and bread wheat B-subgenome, as well as between the bread wheat A- and D-subgenomes and their diploid donors (*T. urartu* and *Ae. tauschii*) along the seven chromosomes based on reduced representative genomic regions. The centromere of each chromosome is highlighted by purple color. Numbers on Y-axis are the values of genetic relatedness. Coordinates of each chromosome are shown in the X-axis.

Next, we estimated the time at which the *Triticum/Aegilops* species diverged from each other based on the same datasets. A genome-wide average was calculated: overall, *Ae. speltoides* diverged from the wheat B-subgenome donor *ca*. 4.49 MYA (95% highest posterior density (HPD): 4.31-4.67 MYA) (**Fig. 2a**). In contrast, bread wheat A- and D-subgenomes diverged from their respective diploid progenitors at much later times: A-subgenome vs. *T. urartu* (1.28 MYA, 95% HPD: 1.22-1.33 MYA), D-subgenome vs. *Ae. tauschii* (<0.88 MYA). Given that wild emmer wheat formed no earlier, and probably much later, than 0.80 MYA^4^, our results rule out the possibility that *Ae. speltoides* is the direct donor to the polyploid wheat B-subgenome. Of the five D-lineage species, our estimates suggest that *Ae. tauschii* evolved independently around 5.37 MYA (95% HPD: 5.16-5.58 MYA), which is probably soon after the homoploid hybridization event between the ancient A- and B-lineages (~5.50 MYA)^4^. The four *Sitopsis* species, *Ae. bicornis*, *Ae. longissima*, *Ae. searsii* and *Ae. sharonensis*, were more recently diversified from a common ancestor <3.73 MYA (95% HPD: 3.58-3.88 MYA). In parallel, we also calculated the divergence times between the seven diploid species and wheat B-subgenome along each of the seven chromosomes (**Supplementary Fig. 4**). We found that *Ae. speltoides* showed later divergence time in centromeric regions than in telomeric regions from the B-subgenome. A similar pattern is observed for the five modern D-lineage species (including the rest four *Sitopsis* species) which also showed variable divergence times along the entire chromosome lengths. In particular, the four D-lineage *Sitopsis* species showed apparent later divergence times from wheat B-subgenome in several subtelomeric regions (*i.e*., chromosomes 2 and 7) than did *Ae. speltoides*. Nonetheless, all the seven diploid species showed earlier divergence times from wheat B-subgenome (>1.00 MYA) than the speciation time of wild emmer wheat (<0.80 MYA) across all seven chromosomes.

Taking advantage of a recently assembled bread wheat cultivar (LongReach Lancer) with its near entire chromosome 2B (*ca*. 450 Mb in length) being substituted by the *T. timopheevii* G-subgenome^27^, together with our assembled sequence contigs of *Ae. mutica* (B-lineage), we constructed a separate phylogeny based on 3,107 RRGRs within this chromosomal segment of all pertinent diploid *Triticum/Aegilops* species and polyploid wheat subgenomes (**Supplementary Fig. 5a**). Overall, we found that the segment-based inferences are highly consistent with the whole-genome molecular phylogenies and divergence times inferred above (**Fig. 2a** and **Supplementary Fig. 3**) and in previous studies^4,21^. For example, *Ae. mutica* belongs to the B-lineage and diverged from *Ae. speltoides* and B-subgenome at a more ancient time (6.37 MYA, 95% HPD: 5.97-6.79 MYA), confirming that *Ae. mutica* can thus be definitely considered as the extant representative most directly related to the B-lineage ancestor^21^. Notably, *Ae. speltoides* diverged from *T. timopheevii* G-subgenome *ca*. 2.85 MYA, (95% HPD: 2.51-3.24 MYA) *i.e*., after its divergence from the B-subgenome progenitor (*ca*. 4.49 MYA). This makes the donor of the G-subgenome substantially older than the estimated allotetraploidization time (<0.4 MYA) leading to speciation of *T. araraticum*, the wild progenitor of *T. timopheevii*^5^. From this analysis, it is clear that *Ae. speltoides* is also not the direct donor to the G-subgenome of *T. timopheevii*, although it is more closely related to the G-subgenome than to the B-subgenome. This is consistent with earlier reports showing that *Ae. speltoides* shares near identical cytoplasmic genomes with *T. timopheevii* (donated by the G-subgenome progenitor) but not *T. turgidum* and *T. aestivum* (donated by the B-subgenome progenitor)^31^. Similar relationships were seen in gel blotting patterns probed by nuclear repeats^32^.

To gain further insights into genome-wide genetic similarities of these *Triticum/Aegilops* species, we calculated genetic relatedness, genetic distance, synonymous (dS) and nonsynonymous (dN) substitution rates based on collinear genes, SCOGs and RRGRs. In line with the divergence times detailed above, the wheat B-subgenome is highly divergent from all the extant diploid *Triticum*/*Aegilops* species, while being most closely related to *Ae. speltoides* (**Fig. 2b** and **Supplementary Fig. 6**). It is notable that the polyploid wheat A- and D-subgenomes display high genetic similarity to their respective diploid donors, *T. urartu* and *Ae. tauschii*, homogeneously across the entire length of each of the seven chromosomes (**Fig. 2c**). However, *Ae. speltoides* shows higher genetic similarity to the wheat B-subgenome in centromeric regions than in telomeric regions (**Fig. 2c**), mirroring the pattern of divergence time detailed above (see **Supplementary Fig. 4**). This bipartite divergence pattern was also observed in the comparisons of the three recently split B-lineage species/subgenomes (*Ae. speltoides*, B- and G-subgenomes) (**Supplementary Fig. 5b**). By contrast, *Ae*. *mutica* (B-lineage) and other *Sitopsis* species (D-lineage) show relatively lower genetic similarities to the wheat B-subgenome (**Fig. 2c** and **Supplementary Fig. 5b**). Together, these genomic features suggest that (*i*) the ancestral B-lineage should have had at least four distinct diploid species, namely *Ae. speltoides*, *Ae*. *mutica*, the progenitors of bread wheat B-subgenome and Timopheevii wheat G-subgenome; (*ii*) the bread wheat B-subgenome is of monophyletic origin, *i.e*., from a single unknown diploid species, now extinct or yet undiscovered, that is phylogenetically close to the extant *Ae. speltoides*; and (*iii*) genetic introgression may have occurred between the diploid progenitor species of bread wheat B-subgenome and other *Aegilops* species.

### Heterogeneous variation pattern and genetic introgression

It has been proposed that interspecific hybridization occurred frequently in many of the *Triticum/Aegilops* species at various evolutionary stages^18,21,33,34^. We thus investigated whether hybridization/introgression also occurred, and if so, to what extent it had shaped the genomes of the five *Sitopsis* species. We found that the B-lineage *Ae. speltoides* and polyploid wheat B-subgenome show distinct phylogenetic topologies in 261 (11.3%) of the 2,314 representative genomic regions, especially at the recombination-active distal chromosome regions (**Supplementary Fig. 7**). Likewise, the five D-lineage species (including the four *Sitopsis* species) also show distinct phylogenetic topologies, but in higher ratio than of the B-lineage, namely in ~35.9% of the total genomic regions (**Supplementary Fig. 7**). The observed heterogeneous patterns along all seven chromosomes suggest the possibilities of either incomplete lineage sorting (ILS) or genetic introgression between the five *Sitopsis* species and their relatives. By computing *D*-statistic, *fd*, hybrid index (*γ*) and *χ* goodness of fit test, we confirm the previously proposed ancestral homoploid hybridization origin of the D-lineage (**Fig. 3a,b** and **Supplementary Fig. 8**) between the ancestral A- and B-lineages^4,21,33,34^.

**Figure 3.**
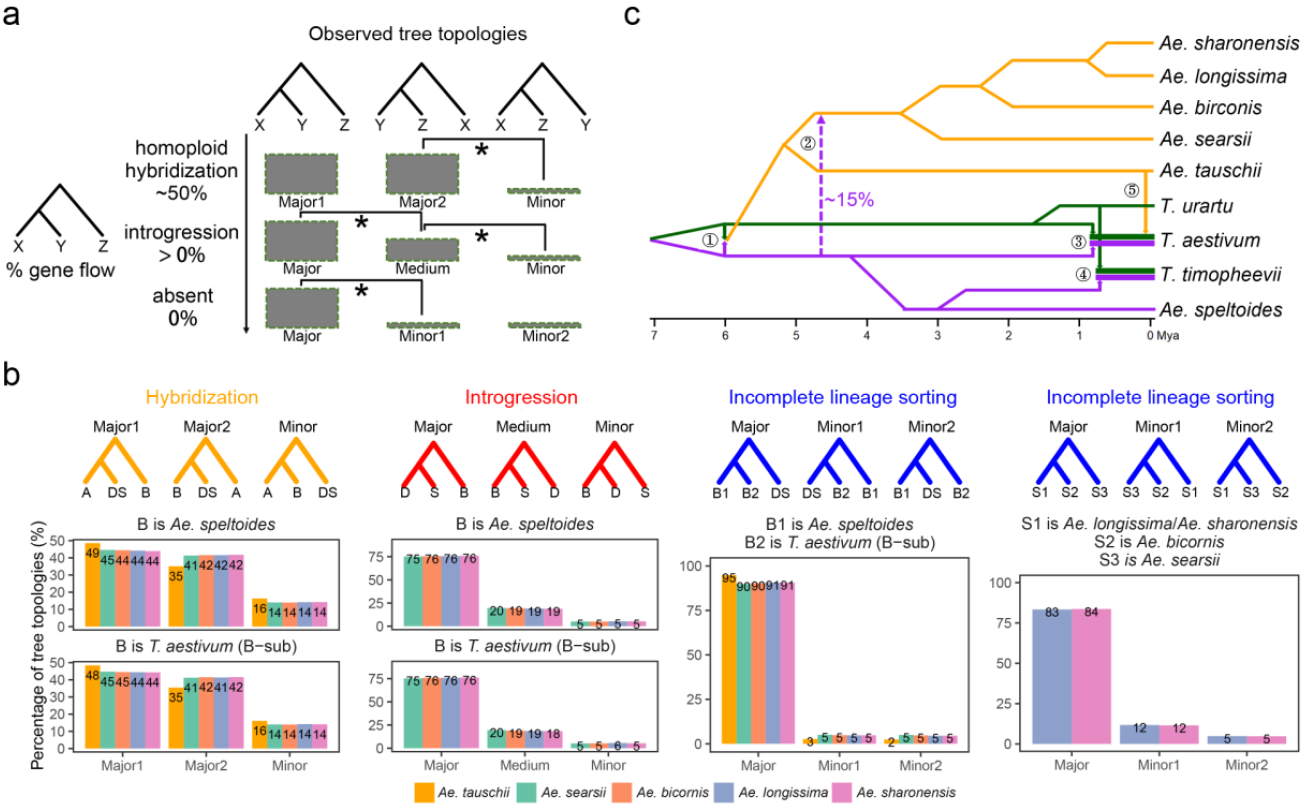
Estimates of genetic introgression in the *Triticum*/*Aegilops* species complex. **a**, Three typical incongruent types between the species tree and gene tree based on reduced representative genomic regions. From up to bottom are: homoploid hybridization, genetic introgression and incomplete lineage sorting (ILS). The three incongruent types are distinguished by the numbers of each tree topology. Significance is determined by the χ^2^-test with *p*-value < 0.001 (shown as asterisk). In homoploid hybridization type, numbers of both the two “major” topologies (“major1” and “major2”, 2 > n_major1_/n_major1_ > 1/2) are significantly higher than the “minor” topology. In genetic introgression type, the number of “major” topology is significantly higher than “medium” topology. Likewise, the medium” topology is also significantly higher than “minor” topology. In ILS type, numbers of both the two “minor” topologies (“minor1” and “minor2”) are less than “major” topology but which are not statistically significant. **b**, Examples of the three incongruent types identified in the *Triticum*/*Aegilops* species complex, including homoploid hybridization between ancestral A- and B-lineages (orange), genetic introgressions from B-lineage (ancestor of *Ae. speltoides* and B-subgenome) to ancestor of the four D-lineage *Sitopsis* species (red), ILS between B- and D-lineages (blue), and ILS among the four D-lineage *Sitopsis* species (blue). Percentages of these trees are shown below the tree topology. Colors in the barplot represent the five D-lineage species. **c**, Evolutionary scenario of the *Triticum*/*Aegilops* species complex based on the integrated estimates of genetic introgression. The numbers ① and ② indicate the ancestral homoploid hybridization between A- and B-lineages and B- to D-lineage ancestral genetic introgression event, respectively. The remaining three numbers (③, ④ and ⑤) represent the allopolyploidization events that formed tetraploid *T. turgidum* ssp. *dicoccoides*, tetraploid *T. araraticum* and hexaploid *T. aestivum*.

Previous cytogenetic studies showed that karyotypes of the four *Sitopsis* species (D-lineage) are more similar to *Ae. speltoides* (B-lineage) than to *Ae. tauschii* (D-lineage)^35^. This observation was also supported by transcriptome-based phylogenetic inference that genetic introgression probably occurred from *Ae. speltoides* to the common ancestor of D-lineage *Sitopsis* species after its separation from *Ae. tauchii*^21^. We propose a different scenario whereby the introgression event more likely occurred from the common ancestor of *Ae. speltoides* and wheat B-subgenome (earlier than 4.49 MYA) to the four D-lineage *Sitopsis* species (**Fig. 3b**). This scenario of ancestral genetic introgression is also confirmed by the distribution pattern of introgressed-sites (*i*-sites) (**Supplementary Fig. 9a**). For example, all the four D-lineage *Sitopsis* species possess relatively more *i-*sites with both the *Ae. speltoides* and bread wheat B-subgenome (1.44% of the total SNPs) (putatively derived from their common ancestor) compared to those of from each of the two B-lineage species (0.90% and 1.04%) and the putative A-lineage (diploid *Triticum*) donor (1.31%). In contrast, the same D-lineage species *Ae. tauschii* harbors similar proportions of *i*-sites to *Ae. speltoides* (0.26%), B-subgenome (0.26%) and their common ancestor (0.30%), but which is markedly lower than the other A-lineage donor, *T. urartu* (0.89%). In line with these observations, allele frequency-based inference of migration also confirmed the ancestral genetic introgression from B- to D-lineage (**Supplementary Fig. 10**). In particular, we identified several genomic regions that show high genetic similarity between the D-lineage *Sitopsis* species and either of the *Ae. speltoides* and bread wheat B-subgenome (**Supplementary Fig. 11** and **12**), suggesting the possibility of genetic introgressions between the B- and D-lineages. These features together may explain why the four D-lineage *Sitopsis* species have higher genetic similarity to the wheat B-subgenome in some genomic regions than to their otherwise phylogenetically closer relative, *Ae. tauschii*. It is notable that previous studies have proposed some additional post-ancestral homoploid hybridization genetic introgressions among the A-, B- and D-lineage species^18,21,33,34^. Broadly consistent with these studies, our integrated analyses also identified genetic introgressions from D- to B-lineage, although some differences were observed between the datasets and methodologies used (**Supplementary Fig. 8** and **10**).

Among the four D-lineage *Sitopsis* species, Waines and Johnson^36^ have proposed that *Ae. sharonensis* is likely a hybrid between *Ae. longissima* and *Ae. bicornis* based on morphology and cytogenetic analyses. Our estimates however did not find evidence for this possibility. The observed heterogeneous pattern was more likely due to incomplete sorting of ancestral polymorphisms (**Fig. 3b**). In line with this conclusion, we found that *Ae. sharonensis* not only possesses high proportion of species-private SNPs (8.80% of the total SNPs) but also shares low proportion of species-shared SNPs with *Ae. bicornis* (1.27%) (**Supplementary Fig. 9b**). It is notable that all the extant D-lineage species were established through a single ancestral homoploid hybridization event, as reported by Marcussen *et al*.^4^ (2014) and modified by Glemin *et al*.^21^ (2019). We thus asked whether the above identified genetic introgressions have differentially sculptured the genomes of the five extant D-lineage species (including the four *Sitopsis* species). Through comparing the distribution patterns of A- and B-lineage specific SNPs, we found that genomic regions containing more A-lineage specific SNPs (A-dominant) are clustered at the recombination-inert proximal regions across all seven chromosomes (**Supplementary Fig. 13**). In contrast, species-specific SNPs identified in either *Ae. speltoides* (B-lineage) or bread wheat B-subgenome (B-lineage) mainly distributed at the recombination-active distal chromosomal regions. In particular, the *Sitopsis* species (excluding *Ae. speltoides*) harbor more B-lineage species-specific SNPs compared to *Ae. tauschii* (D-lineage), confirming the above identified B- to D-lineage introgression in the *Sitopsis* species. Together, our genome-scale estimates revealed frequent post-ancestral homoploid hybridization introgressions among the *Triticum*/*Aegilops* species (**Figure 3c**), which may have shaped the genomes of extant *Sitopsis* species.

### Post-speciation amplification of transposable elements

The genome features detailed above revealed differences in genome content between the five *Sitopsis* species and their close relatives (see in **Figure 1** and **Table 1**). We thus assessed whether the differences in genome content are due to the genetic introgressions (see in **Figure 3**) or independent expansion/contraction of transposable elements (TEs). At the overall level, our analyses revealed that 2.73-3.95 Gb (77.6%-81.2%) of the genome components of these *Triticum/Aegilops* species are composed of the *gypsy-like, copia-like* and *CACTA* TE families (**Supplementary Fig. 14a-b**). Between the two B-lineage species/subgenome, about 0.97 Gb (90.4%) of the genome size difference can be attributed to the high copy numbers of *gypsy-*like and *CACTA* TEs in polyploid wheat B-subgenome relative to *Ae. speltoides* (**Supplementary Fig. 14c**). In the D-lineage, compared to *Ae. tauschii*, all three types of TEs (*gypsy-like, copia-like* and *CACTA*) show higher copy abundance in the two earlier established species *Ae. bicornis* (1.39 Gb, accounting for 82.9%% of the genome size difference) and *Ae. searsii* (1.00 Gb, 89.8% of the difference). In contrast, only *gypsy-like* and *copia-like* TEs exhibit high copy numbers in the two more recently established *Sitopsis* species, *Ae. longissima* (1.47 Gb, 93.6% of the difference) and *Ae. sharonensis* (1.54 Gb, 92.3% of the difference).

To examine whether specific repetitive sequences have contributed to the differences in genome content, we further characterized 85 retrotransposon and transposon subfamilies that are responsible for >90.0% of the genome size differences among the *Triticum/Aegilops* species (**Fig. 4a**). In line with the above results, differential abundance of two *gypsy-like* subfamilies (*RLG_famc3.1* and *RLG_famc3.4*) account for about 10.7% of the genome size differences between the polyploid wheat B-subgenome and *Ae. speltoides* (**Fig 4a**). Likewise, different copy numbers of 29 subfamilies are responsible for the different genome content among the five D-lineage species. Intersection analysis showed that the 20 and 19 lineage-specific retrotransposon and transposon subfamilies contributed to 625.8 Mb (58.5% of B-lineage) and 479.4-1027.4 Mb (43.0-63.0% of D-lineage) of the genome size differences in the B- and D-lineage species, respectively (**Fig. 4b** and **Supplementary Table 1**). In contrast, seven subfamilies that were shared between the B and D-lineages account for 383.2 Mb (35.8% of B-lineage) and 466.7-502.8 Mb (29.5-42.0% of D-lineage) of the differences in genome contents. It suggests that genome size differences within the B- and D-lineages are primarily due to distinct proportions of the specific TE subfamilies.

**Figure 4.**
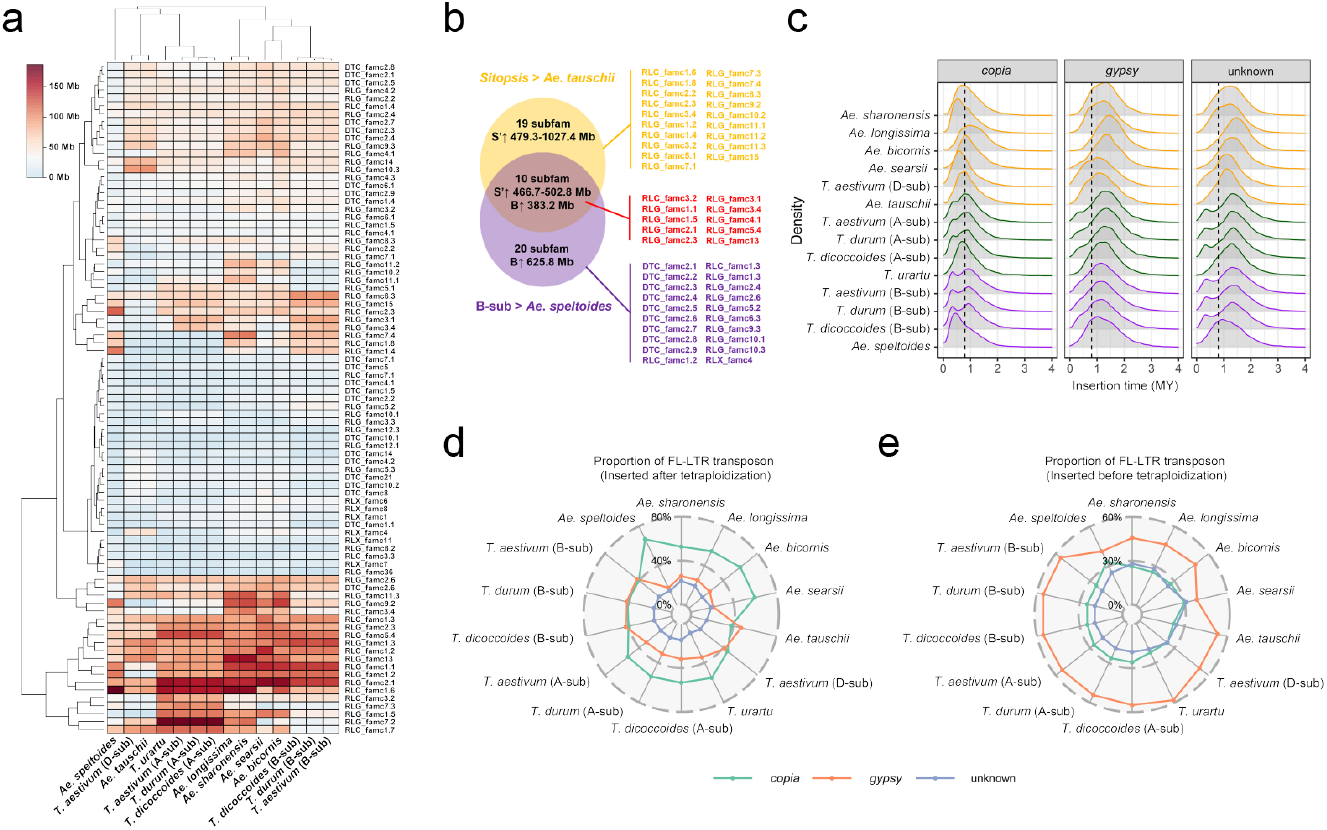
Evolutionary dynamics and insertion times of transposable elements in the *Triticum/Aegilops* species complex. **a**, Heatmap of the lengths of the 85 transposable element (TE) subfamilies in the diploid species and polyploid wheat subgenomes. Scale bar on the top left corner denotes the length of each TE subfamily. **b**, Intersection analysis of the TE subfamilies that are expanded specifically in B- (purple color, 20 subfamilies) and D-lineages (orange color, 19 subfamilies), as well as these shared between the two lineages (red color, 10 subfamilies). Total length of these expanded TE subfamilies are shown in each section. The character S’ and B indicate the four D-lineage *Sitopsis* species and bread wheat B-subgenome, respectively. Black arrows after the S’ and B indicate the increased genome size in these genomes compared to their close relatives. The expanded TE subfamilies are listed on the right. **c**, Insertion times of the retrotransposons in the diploid species and polyploid wheat subgenomes. Green, purple and orange colors represent the A-, B- and D-lineages, respectively. **d** and **e**, Proportions of the expanded full length long-terminal repeat (FL-LTR) transposons in the diploid species and polyploid wheat subgenomes before and after the tetraploidization speciation event (~0.8 MYA). Light green, orange and blue colors are the *copia*-like, *gypsy*-like and unknown retrotransposons, respectively.

We next estimated the burst time of retrotransposons to reexamine whether they were expanded or contracted independently in the B- and D-lineages. If the retrotransposons expanded in the B- and D-lineages are directly derived from the above identified inter-specific genetic introgressions (detailed in **Fig. 3b**), we would expect to identify pre-introgression (>4.49 MYA) burst of the retrotransposons in the two lineages. However, our estimates identified relatively recent retrotransposon amplifications (<3.00 MYA) in all the diploid species and polyploid wheat subgenomes (**Fig. 4c**). It is notable that both the A- and B-subgenomes of the three polyploid wheats (emmer, durum and bread) have experienced a common recent retrotransposon expansion (~0.5 MYA), most likely after the allotetraploidization event *ca*. 0.8 MYA. Consistent with this inference, their diploid donors (*T. urartu* and *Ae. tauschii*) and the five *Sitopsis* species do not share this recent retrotransposon burst. We next compared the insertion times of these retrotransposon families relative to the allotetraploidization event (0.8 MYA). In the B-lineage, about 55.6-55.7% of the earlier expanded retrotransposons (>0.8 MYA) in polyploid wheat B-subgenome can be attributed to the *gypsy*-like retrotransposons (**Fig. 4 d-e** and **Supplementary Table 2**). However, *Ae. speltoides* possesses more recently (<0.8 MYA) amplified *copia*-like retrotransposons (67.0% of the total) compared to the polyploid wheat B-subgenome (40.5%-41.2%). It may explain why *Ae. speltoides* shows distinct distribution density of *copia-like* retrotransposons compared to B-subgenome and the other *Sitopsis* species (see in **Figure 1**). In the D-lineage, the *copia-like* families are responsible for 52.6-59.9% of the recent amplified retrotransposons (<0.8 MYA) in the four *Sitopsis* species (**Fig. 4 d-e** and **Supplementary Table 2**). In contrast, only 37.8-45.1% of the recent expanded retrotransposons are *copia*-like families in polyploid wheat D-subgenome and its donor *Ae. tauschii*, supporting the above observed distinct evolutionary histories of the five modern D-lineage species. In the A-lineage, slightly lower proportions of recently expanded *copia*-like families were identified in the polyploid wheat A-subgenome (31.2-31.9%) compared to their diploid donor *T. urartu* (34.6%). We also estimated the insertion times for the expansion of the above identified TE subfamilies in the seven diploid species and wheat B-subgenome. All these TE subfamilies possess insertion times <3.00 MYA (**Supplementary Fig. 15**), which are later than the ancestral B- to D-lineage introgression (4.49 MYA) (detailed in **Fig. 2a**). In particular, all the four D-lineage *Sitopsis* species possess distinct expansion patterns of these TE subfamilies compared to *Ae. speltoides* and B-subgenome, even those that are expanded in both the B- and D-lineages (**Supplementary Fig. 15**). As these retrotransposon and transposon subfamilies are responsible for >90% genome size differences, it suggests that the increased genome content in D-lineage *Sitopsis* species compared to *Ae. tauschii* is more likely due to the post-speciation expansions/contractions of a few specific active TEs rather than to the direct B- to D-lineage genetic introgression.

### Pan-genomic analyses of the Triticum/Aegilops species

Pan-genomic analyses of the *Triticum/Aegilops* species were performed based on protein-coding genes and genome structural variations (SVs). The five *Sitopsis* species contain 23,456-24,344 gene families, 56.3% (17,595) of which are shared with the other diploid and polyploid wheat species, probably representing the core gene set of *Triticum*/*Aegilops* species complex (**Fig. 5a**). In addition, a total of 11,580 (34.4%) dispensable and 2,798 (9.3%) species-specific gene families were also identified from these *Triticum/Aegilops* species. Of these orthologous gene families, from 419 (1.11%) to 1,086 (2.56%) species-specific genes and from 1,406 (3.78%) to 1,455 (3.79%) specifically expanded genes were identified in the five *Sitopsis* species (**Fig. 5a**). Functional analyses of the *Sitopsis*-specific and expanded genes reveal significant enrichment in basic cellular activities, including DNA recombination, DNA integration and metabolic process (**Fig. 5b** and **Supplementary Table 3**). Based on the same protein-coding gene set, we characterized evolutionarily conserved genomic regions by identifying the shared syntenic orthologous genes in the *Triticum*/*Aegilops* species. Our results reveal that centromeric regions of all seven chromosomes possess very few numbers of core putative proto-genes (pPGs) (present in all diploid species and polyploid wheat subgenomes) (**Supplementary Fig. 16**), suggesting the low level of genetic conservation of centromeric regions in the *Triticum*/*Aegilops* species. In addition, we identified several large genomic regions that are evolutionarily non-conserved in the B-and D-lineages. For example, two large non-conserved genomic regions near the telomere of chromosome 2 are potentially correlated with the evolutionary divergence between the five *Sitopsis* and their close relatives, wheat B-subgenome (B-lineage) and *Ae. tauschii* (D-lineage) (**Supplementary Fig. 16c** and **e**).

**Figure 5.**
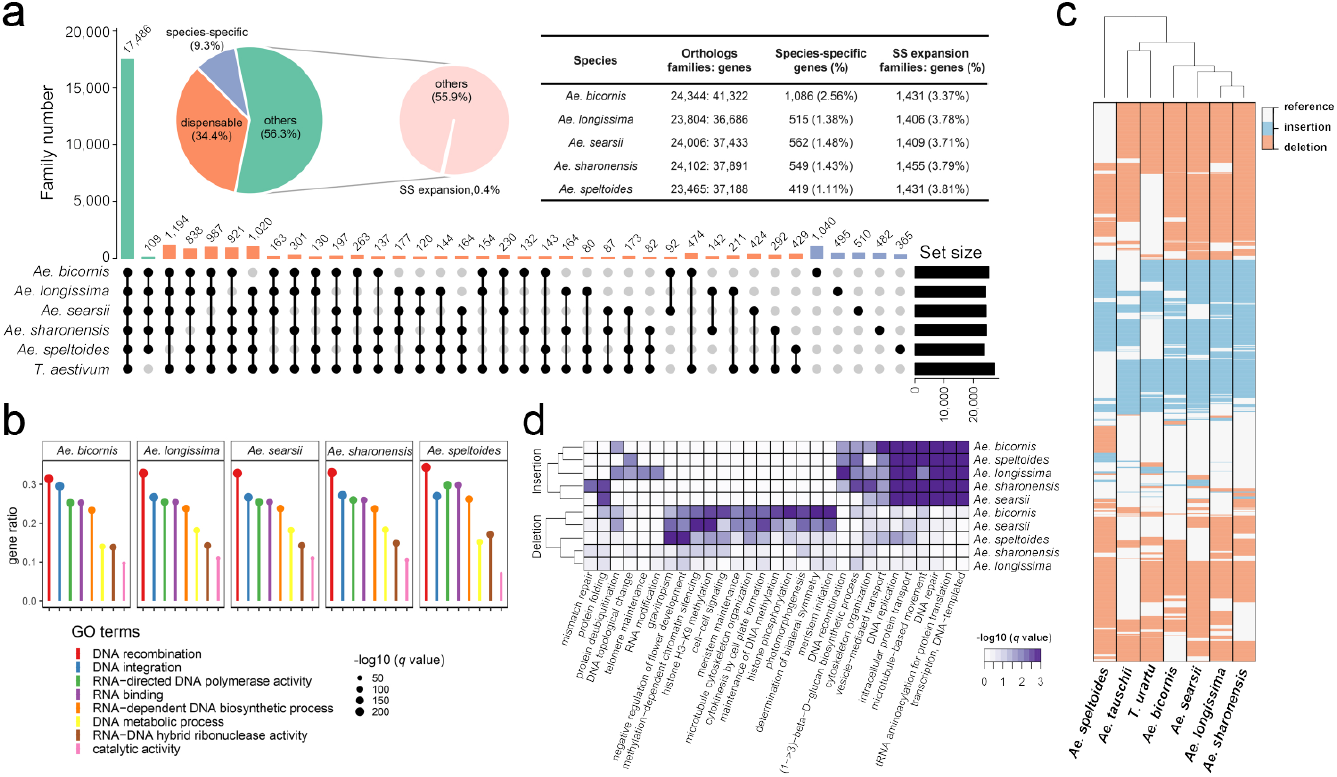
Pangenomic analyses of the *Triticum/Aegilops* species based on protein-coding genes and genome structural variations. **a**, Intersection analysis of the protein-coding genes in the *Triticum/Aegilops* species. All the polyploid wheat subgenomes and their diploid donors are treated as a single gene set. Light green, orange and blue colors are the core (others), dispensable and species-specific gene families in the *Triticum/Aegilops* species complex. SS expansion gene families (0.4% of the total) are defined as present in all the *Triticum/Aegilops* species (the 56.3% core gene families) but expanded specifically in the five *Sitopsis* species. Numbers in the up-corner table indicate the gene numbers of *Sitopsis*-specific and SS expansion gene families. **b**, Functional enrichment analysis of the SS expansion genes in the *Sitopsis* species. **c**, Heatmap of the distribution of genome structural variations in the seven diploid species compared to bread wheat B-subgenome. Orange, blue and white colors represent deletion, insertion and the same as bread wheat B-subgenome. **d**, Functional annotations of the SV-related genes in the five *Sitopsis* species. Full list of the GO terms is shown in Supplementary Table 5.

Pan-genomic analyses were also performed with the genome-wide SVs characterized from the seven diploid *Triticum/Aegilops* species. A total of 38,994 common SVs were identified in the five *Sitopsis* species (ranging from 37,039 to 37,721 in each species), 18,153 of which are shared with the two diploid species, *Ae. tauschii* (DD) and *T. urartu* (AA) (**Supplementary Fig. 17a**). Of the shared SVs, 16,337 monomorphic SVs that are fixed in the seven diploid species compared to polyploid wheat B-subgenome were excluded, leaving 1,816 polymorphic insertions/deletions in the genome-scale SV dataset. Further analyses of the 1,816 polymorphic SVs show that while these SVs scattered randomly along the seven chromosomes (**Supplementary Fig. 18**), the seven diploid species possessed five to 120 species-specific SVs (**Fig. 5c** and **Supplementary Fig. 17b-c**). For example, while *Ae. speltoides* is phylogenetically close to the B-subgenome, it still carries 120 (6.61% of the total polymorphic SVs) species-specific SVs compared to the B-subgenome and all the other diploid species. Compared to the above *Sitopsis*-specific and expanded genes that are mainly involved in basic cellular activities, the SV-associated genes identified in the five *Sitopsis* species are correlated with several functional important phenotypes, such as photomorphogenesis, DNA methylation, chromatin silencing and topology, meristem and flower development (**Fig. 5d** and **Supplementary Table 4**).

### The Sitopsis genomic resource

The *Aegilops* species represent the secondary gene and germplasm pools for wheat genetic improvement^37^. We thus examined whether the five *Sitopsis* species contain homoeologous genes related to agronomic and pathogen-resistant traits of durum and bread wheats. Our genome-wide screening of the functional nucleotide-binding site and leucine-rich repeat (NBS-LRR) domains identified a total of 5,867 genes in the 14 diploid species and polyploid wheat subgenomes. Further redundancy analysis revealed that the total NBS-LRR gene pool consists of 2,573 (95% identity) to 4,439 (100% identity) unique genes in the *Triticum/Aegilops* species, with 90% of the NBS-LRR genes being contained in 12 (95% sequence identity) and 13 (100% sequence identity) wheat genomes, respectively (**Supplementary Fig. 19a**). It suggests that the 5,867 NBS-LRR genes may represent the core resistant gene set of *Triticum/Aegilops* species. In particular, the 5,867 NBS-LRR genes are mostly distributed at the distal chromosomal regions in all the *Triticum/Aegilops* species (**Supplementary Fig. 19b**). This is broadly consistent with previous findings that gene families and QTLs associated with adaptation to biotic and abiotic stresses are mainly clustered near the subtelomeric chromosome regions in polyploid wheat^25–27^.

We noted that the five *Sitopsis* species (388-490) contain relatively more NBS-LRR gene families than do the other two diploid species, *Ae. tauschii* (350) and *T. urartu* (318), and the two subgenomes of wild emmer wheat (239-286), but are comparable to the domesticated durum (326-435) and bread wheat (401-542) (**Supplementary Fig. 20a**). Intersection analysis of these NBS-LRR gene families allocated 116 (38.8%), 167 (57.5%) and 12 (3.7%) as core, dispensable and *Sitopsis*-specific gene families (**Supplementary Fig. 20b**). The 116 core and 167 dispensable NBS-LRR gene families are the major components of innate immune system in the *Triticum/Aegilops* species. Thus, the 11 *Sitopsis*-specific NBS-LRR gene families may provide specific genetic resources for the improvement of disease resistance in domesticated durum and bread wheats. In addition, we also characterized a set of homoeologous genes in the five *Sitopsis* species, which are related to stripe rust (*Puccinia striiformis* f. sp. *tritici*, *Pst*) and powdery mildew (*Blumeria graminis* f. sp. *tritici*, *Bgt*) resistance (**Supplementary Table 5**). Because *Pst* and *Bgt* are two major fungal diseases causing heavy yield loss of wheat worldwide^38^, the novel resistant genes we identified in the *Sitopsis* species might be important genetic resources for future wheat breeding.

For agronomic traits, we checked the copy number and nucleotide variation pattern of two major domestication genes related to the non-shattering phenotype, namely, free-threshing seed (*Q/q*) and nonfragile rachis (*Btr/btr*) (**Supplementary Table 5**). The *Q*/*q* gene encodes an *AP2*-like transcription factor that confers free-threshing and also has pleiotropic effects on a number of other domestication traits, including rachis fragility, spike architecture, and flowering time^39–41^. The seven diploid *Triticum/Aegilops* species and their attendant natural and resynthesized polyploids with diverse genome combinations, including BBAA, S^sh^S^sh^A^m^A^m^, S^l^S^l^AA, S^b^S^b^DD and AADD, all show substantial morphological differences in inflorescence structure^42^ and distinct *Q/q* allele expression patterns^43^. Here we show that while all the five *Sitopsis* species harbor the wild type *q* allele (L329), it is different from that of the durum and bread wheat A-subgenome domesticated *Q* allele (I329) and their diploid donor *T. urartu q* allele (V329), and contains numerous unique synonymous and nonsynonymous mutations (**Supplementary Fig. 21** and **Dataset**). Similar phenomenon was also observed in the two nonfragile rachis genes (*Btr1* and *Btr2*) in which many genetic variants are found in the five *Sitopsis* species (**Supplementary Table 5** and **Dataset**). It has been documented that novel SNPs in the miRNA binding site at *Q*/*q* gene are correlated with changes in transcriptional regulation and plays pleiotropic roles in growth and reproductive development^44^. Thus, the natural variations we identified in the *Sitopsis* species are potentially valuable in breeding new wheat cultivars. In addition, we also characterized candidate genes that are functionally associated with other important agronomic traits (*i.e*., tiller number and kernel size) and floral development (*i.e*., vernalization and photoperiod-insensitive) (**Supplementary Table 5**). These genic and genetic resources might be key variants for future wheat breeding or *de novo* domestication of new types of wheat.

## Discussion

We have assembled chromosomal level reference genomes of all five *Aegilops* species of the *Sitopsis* section and conducted comparative genomic analyses both among the five species and with the other available diploid species and polyploid wheat subgenomes. Our main motivation was to better understand the evolutionary histories and trajectories of the *Sitopsis* species and especially, the origin of the polyploid wheat B-subgenome as it has long been debated. A long-standing hypothesis posited that the wheat B-subgenome was derived monophyletically from *Ae. speltoides*^9^. This was formulated based on multiple lines of observational and empirical evidence, including botanical, cytological, phylogenetic and biogeographical^37^. Although this hypothesis was already questioned nearly 50 years ago based on the near absence of homologous synapsis between the S- and B-subgenome chromosomes in artificial hybrids involving both higher- and lower-pairing types of *Ae. speltoides*^10,11,20^, it was revised by an extensive molecular marker-based population study^16^. An alternative hypothesis is that both the extant *Ae. speltoides* and wheat B-subgenome/its progenitor diverged from their original common ancestor at both the diploid and polyploid levels^8,12,17,45^. Our genome-scale comparative analyses show that *Ae. speltoides* and the B-subgenome have diverged ~4.49 MYA, *i.e*., at a much earlier time than the speciation of tetraploid emmer wheat approximately 0.8 Mya^4^. In other words, major divergence between *Ae. speltoides* and the B-subgenome diploid donor should have occurred at the diploid level. Moreover, the estimates of genome-wide genetic similarity between B-subgenome and *Ae. speltoides* is far less than those of the A- and D-subgenomes from their respective diploid donors, *T. urartu* and *Ae. tauschii*. Together, our results lead to the unequivocal conclusion that *Ae. speltoides* is not the direct progenitor to the B-subgenome. Similarly, based on an independent analysis of a chromosomal segment corresponding to a *ca*. 450 MB segment covering almost the entire chromosome 2B^27^, we show that *Ae. speltoides* is also not the direct donor to the G-subgenome of *T. timopheevii*.

Another hypothesis for the origin of the wheat B-subgenome poises that it had formed through multiple hybridizations and introgressions of diverse genomic sequences from *Sitopsis* species at the tetraploid level. According to this polyphyletic scenario, the tetraploid wild emmer wheat (*T. turgidum* ssp. *dicoccoides*) was likely established through the intercrossing of two or more amphiploids with the same A genome species (*T. urartu*) but different S-genome donors (*Sitopsis* species)^8,19^. However, the observed heterogeneous genomic patterns between the five *Sitopsis* species and wheat B-subgenome are more likely due to incomplete sorting of ancestral alleles and B-to-D lineage genetic introgressions. This refutes polyphyletic origins of wheat B-subgenome from diverse *Sitopsis* species at the tetraploid level. Alternatively, the direct progenitor of wheat B-subgenome itself might be of polyphyletic origin through the hybridizations/introgressions between two or more distant ancestral B-lineage species at the diploid level^18^. However, our comparisons clearly show that the four extant B-lineage species/subgenomes (*Ae. speltoides*, *Ae. mutica*, B-subgenome and G-subgenome) are evolutionarily independent from each other. Together, we conclude that the direct donor of the B-subgenome is a distinct diploid species that diverged from *Ae. speltoides* 4.49 MYA, but which experienced genetic introgressions with the D-lineage *Sitopsis* species before its hybridization with *T. urartu* leading to formation of *T. turgidum*. However, it still remains mysterious why both of the diploid progenitor species to the B- and G-subgenomes went extinct while their two congeneric species, *Ae. speltoides* and *Ae. mutica*, are extant. It might be that both diploid donors were out-competed by their tetraploid progeny species, *T. turgidum*, ssp. *dicoccoides* and *T. araraticum*, the wild progenitor of *T. timopheevii*, or that the B and G donors remain to be discovered. The latter is possible but not very likely considering that there has been much effort to find these species in the levant.

We also performed genomic comparisons to elucidate evolutionary dynamics of the *Triticum*/*Aegilops* species complex. Our results reveal high collinear genome structure among the *Sitopsis* species, albeit they all contain high but markedly variable proportions of repetitive sequences. The differences in genome size among the *Triticum*/*Aegilops* species are primarily due to independent post-speciation amplification of a few specific TEs. In addition, we show how detailed comparisons between the reference-quality genome assemblies of the *Sitopsis* species and the wheat subgenomes may open new avenues for the utilization of this set of important genic and genetic resources (*i.e*., homoeologous genes related to agronomic and pathogen-resistant traits) for future wheat breeding. The high-quality genome assemblies for the *Sitopsis* species together with those of other species in the *Aegilops*/*Triticum* complex enable an unprecedented opportunity for further evolutionary, genetic and breeding studies in the wheat group.

## Acknowledgements

We thank Moshe Feldman for critical reading and constructive comments. This study was supported by the Natural Science Foundation of China (#31991211 to B.L. and #31970235 to L.F.L.), the Shanghai Pujiang Program (#19PJ1401500 to L.F.L.), the Israel Science Foundation (ISF)-China National Natural Science Foundation (NSFC) collaborative grant to B.L (#32061143001) and A.A.L. (#3394/20) and China Postdoctoral Science Foundation Grant (2021M690683).

## Additional information

Supplementary information is available for this manuscript at xxx.

## Author contributions

L.F.L., A.A.L., and B.L. conceived this project and coordinated research activities; Y.S. and Y.L. collected and maintained the plant materials; Y.S., Y.W. and J.Z. took the photos of the spikes of the *Triticum/Aegilops* species and carried out the FISH experiments; Y.W. performed the flow cytometry; Z.H.W. conducted the genome collinear comparisons; F.M. Z.B.Z. and X.F.W. reconstructed the phylogeny and estimated the divergence time and genetic introgression; Z.H.W., N.L. and Z.B.Z. carried out the pan-genomic analyses; N.L. and Z.B.Z. annotated the repetitive elements; N.L. analyzed the genome structural variations; Z.B.Z., Y.S. and Z.H.W. identified the domestication and resistance genes; N.D. and X.F.W. carried out the synonymous and non-synonymous substitution analyses; L.F.L., A.A.L., G.L., F.M. and B.L. wrote the manuscript. All authors discussed the results and approved the manuscript.

## Online content

Any methods, additional references, Nature Research reporting summaries, source data, statements of data availability and associated accession codes are available at xxxxx. All sequence reads assemblies have been deposited into the National Center for Biotechnology Information under accession no. PRJNA700474.

## Online Methods

### Plant materials, DNA and RNA extraction

All the five *Sitopsis* species were grown in greenhouse under following conditions: temperatures of 25/16 °C (day/night), photoperiod of 16/8 hour (day/night). Specimen identification was performed by checking the spike morphology and chromosome karyotype. Inflorescence morphology was captured by digital single lens reflex camera (EOS 6D MARK II) in photo studio. Karyotypes of the five species were checked using fluorescence in situ hybridization (FISH) according to Han *et al*. (2005)^46^ and Kato *et al*. (2004)^47^ with minor modifications. The haploid genome size was estimated with flow cytometry using Attune focusing analyzer (ABI, CA, USA). Genomic DNA was isolated from fresh leaf tissue using CTAB. Total RNAs were extracted from root, leaf and inflorescence tissues independently based on standard TRIzol^@^ protocol (TRIzol).

### Genome assembly, gene annotation and quality assessment

Chromosomal level reference genomes of the five *Sitopsis* species were assembled by a combination of Oxford Nanopore Technologies (ONT) single-molecule real-time technology and Hi-C based scaffolding strategy, followed by Illumina short read-based polishing. In brief, about 740-799 Gb (~114-178× genome coverage) high quality ONT long reads were generated using the PromethION platform and 262-302 Gb (~39.8-51.4×) short reads were obtained from the Illumina Noveseq platform (**Supplementary Table 5**). The long ONT reads were corrected using Canu^48^ and assembled to long contigs by wtdbg2^49^. Then, the draft assemblies were polished using Racon^50^ and Pilon^51^ separately. The Hi-C data (~183-256×) was used to link the polished contigs into seven pseudochromosomes using LACHESIS^52^.

Protein coding genes and non-coding RNAs were predicted using *de novo* and homology gene blast strategies (see details in **Supplementary Notes**). All predicted protein coding genes were annotated based on KOG, KEGG and GO databases. Repetitive elements were identified by LTR_FINDER^53^ and RepeatScout^54^, and then annotated using RepeatMasker (http://www.repeatmasker.org). Quality validation of the genome assemblies were assessed by estimating the completeness of the gene repertoire using CEGMA^55^ and BUSCO^56^.

### Phylogeny, divergence time and genetic introgression

Phylogenetic topologies and divergence times of the *Triticum*/*Aegilops* species and outgroups were estimated based on whole genome resequencing data and single copy orthologous gene. Genome sequences and resequencing data of the five *Sitopsis* species were generated in this study. The other *Triticum*/*Aegilops* and outgroup species were obtained from previously released reference genomes (**Supplementary Notes**). Phylogenetic trees were constructed using maximum likelihood method implemented in RAxML (v.8.2.12)^57^ with GTR-GAMMA substitution model and 1,000 bootstrap replicates. Divergence times of the *Triticum*/*Aegilops* species were estimated using the program BEAST (v.2.6.0)^58^. Tree topologies were summarized using TreeAnnotator (v.2.6.0)^58^ and visualized by ggtree^59^.

Genome-wide inter-specific genetic divergence was evaluated for the *Triticum*/*Aegilops* species by calculating the synonymous (dS), non-synonymous (dN) mutation rate based on PAML^60^. In addition, we also estimated pair-wise branch length and genetic similarity using *DendroPy* module^61^ in Python. Genetic introgression among the A-, B- and D-lineages were estimated using a combination of the D-statistic, fd, hybrid index (γ) and χ^2^ good ness of fit test^62–64^. Introgressed variants among the four D-lineage *Sitopsis* species were identified based on the shared-specific SNPs for each species pair compared to the other species.

### Repetitive elements classification and expansion

Full-length LTR retrotransposons in the five *Sitopsis* and the other *Triticum*/*Aegilops* species were characterized using LTR-harvest^65^ and LTR-finder^53^. Then, the program RepeatScout^54^ was employed to build consensus LTR retrotransposon sequences. DNA transposons were identified by a homology search against the REdat_9.7_Triticeae subset of the PGSB transposon library^66^ using vmatch (http://www.vmatch.de). The above identified DNA transposons and LTR retrotransposons were classified into different families using RepeatMasker (http://www.repeatmasker.org) and CLARI-TE procedures^67^. Insertion time of each LTR retrotransposon family was estimated using the formula: age = K/2r, where K is the Kimura 2-parameter distance and r is the mutation rate of 1.3×10^-8 28^. K-mer analyses of these wheat genomes were performed by KAT^68^ for each genome/subgenome.

### Genome collinearity and pan-genomic analyses

Genome collinearity between the five *Sitopsis* and the other *Triticum/Aegilops* species was performed using MCscanX^69^. Inter-specific homologous genes were characterized using BLASTP with the default parameters. The resulting syntenic genomic regions were subjected to identify orthologous genes using ColinearScan^70^. Putative protogene (pPG) was characterized for the A-, B- and D-lineage respectively according to previously published protocols^71,72^.

Pan-genomic analyses were performed based on both the protein coding genes and genome structural variations (SVs). For the protein coding genes, all the three polyploid wheat subgenomes (A, B and D) and their diploid donors (*Ae. tauschii* and *T. urartu*) were mixed as a new *in silico* species called “ABD”. Then, the five *Sitopsis* species and “ABD” were used to identify gene families using OrthoFinder^73^. Gene family expansion and contraction were inferred using previous established phylogenomic approach^25^. Log-transformed gene family size among fifteen genomes and/or subgenomes were compared using the phylANOVA function of phytools package (https://github.com/liamrevell/phytools) in R according to the guidance of the phylogenetic species tree. Genome SVs of the *Ae. tauschii, T. urartu* and five *Sitopsis* species were characterized based on the ONT data. The corrected ONT long reads were mapped onto the bread wheat (Chinese Spring) B-subgenome IWGSC_v1.0 using minimap2^74^ and predicted using Sniffles^75^. Only the unique mapped reads with mapping quality >30, depth >10 and alternative allele ratio >0.2 were kept for subsequent analyses. GO enrichment of the candidate protein coding and SV-related genes were performed using ClusterProfiler^76^.

### Identification of resistance and agronomic-related gene

The nucleotide-binding and leucine-rich repeat immune receptor (NLR) genes were identified using NLR-Annotator pipeline^77^. In brief, candidate NLR genes were predicted by searching CDS sequences against the Pfam database (https://pfam.xfam.org). Custom Python script was used to classify NLR genes according to the position of intron located inside or outside of NB_ARC domain. The other agronomic-related genes in the five *Sitopsis* species were identified by searching the CDS sequences of candidate genes onto the genome assemblies using BLASTN with E-value < 10^e-5^ and hits length > 300bp. All candidate hits were manually checked and compared with corresponding protein annotation databases.

