## Supplementary notes and figures for "Genome sequences of the five *Sitopsis* species of *Aegilops* and the origin of polyploid wheat B-subgenome"

**Address**:
[1] Key Laboratory of Molecular Epigenetics of the Ministry of Education (MOE), Northeast Normal University, Changchun 130024, China;
[2] Ministry of Education Key Laboratory for Biodiversity Science and Ecological Engineering, School of Life Sciences, Fudan University, Shanghai 200438, China;
[3] Department of Plant and Environmental Sciences, The Weizmann Institute of Science, 76100 Rehovot, Israel;
[4] Faculty of Agronomy, Jilin Agricultural University, Changchun 130118, China.

**Supplementary information**

**This file includes:**

Supplementary Notes

Supplementary Figures 1-23

### Supplementary Notes

#### ***Genome survey, assembly and Hi-C scaffolding***

Genome features of the five *Sitopsis* species were surveyed by GenomeScope[78] based on Illumina short reads. The Illumina DNA paired-end libraries were constructed with an insert length of 350 bp, and 262.71-302.73 Gb (44.53-66.42× coverage) sequencing data were produced on the Illumina Novaseq platform (Illumina, San Diego, CA, USA) according to the manufacturer's instructions (**Supplementary Fig. 22**). Genome size and heterozygosity of the five species varied from 4.49-6.60 Gb and 0.01-0.37, respectively. Reference genomes of the five species were assembled by a combination of short-read Illumina and long-range Nanopore sequences. DNA libraries for real-time single-molecule sequencing were prepared according to the manufacturer's instructions, and sequenced on Nanopore PromethION sequencer (Oxford Nanopore Technologies, Oxford, UK). About 739.62-798.86 Gb long reads were obtained from the five species, with contig N50 length of 28,616-34,991 bp and mean length of 23,221-27,320 bp (**Supplementary Table 6**). The long sequence consensus was corrected using Canu[48] and assembled using Wtdbg2[49] and NextDenovo (https://github.com/Nextomics /NextDenovo). Genome assemblies of the five species were polished by Racon[50] and Pilon[51] based on long-read and short-read data, respectively. The draft assembled genomes consist of 1,154-35,663 contigs, with contig N50 length of 563,410-8,720,942 bp (**Supplementary Table 7**). The genome completeness was assessed using a terrestrial plant dataset from BUSCO[56] and a eukaryote dataset from CEGMA[55] (Parra *et al*., 2007). All the five *Sitopsis* species show high level of genome completeness (BUSCOs: 92.92-95.10%; CEGs: 95.85-99.13%) (**Supplementary Table 8**). Chromosomal level reference genomes were generated by link the contigs into seven pseudochromosomes based on the Hi-C data using LACHESIS[52]. About 1,051.32-1,234.93 Gb (179.15-280.09x genome coverage) Hi-C short reads were generated for the five species on Illumina Novaseq platform (Illumina, San Diego, CA, USA) (**Supplementary Table 9**). The construction of pseudomolecules was validated by counting the valid interaction read pairs within each 500-kb bin. The heatmap revealed high interactions between the adjacent genomic regions of all the seven pseudomolecules (**Supplementary Fig. 23**). A total of 49.06-57.68% (95.21-97.07% of the assembled genome) of the contigs were clustered, 47.55-53.62% (90.83-93.44% of the genome) of which were ordered on the on the seven pseudomolecules (**Supplementary Table 10**).

#### ***Annotation of gene and repetitive sequence***

Protein-coding genes were annotated by the combination of *de novo*, homeolog and unigene prediction strategies. The *de novo* annotation of the five species was performed using the programs Genscan[79], Augustus[80], GlimmerHMM[81], GeneID[82] and SNAP[83]. Gene homeolog was predicted using GeMoMa[84]. Unigenes were identified based on the transcriptome data generated from the leaf and root tissues of each species. The transcriptome data were mapped on the assembled genomes using Hisat2[85] and Stringtie[86]. Then, the programs TransDecoder (https://github.com/TransDecoder/TransDecoder) and GeneMarkS-T[87] were employed to predict the protein-coding genes. In addition, we also used the program PASA[88] to predict protein-coding genes. All the protein-coding genes annotated by the above three strategies were combined by EVM[89]. About 61,084-62,804 high confidence genes were predicted in the five species (**Supplementary Table 11**). The total length of the protein-coding genes varied from 213,009,021-319,640,000 bp, with each gene being on average 3,364-5,168 bp long in the five species. Functions of these protein-coding gene were annotated by searching against the databases NR (https://www.ncbi.nlm.nih.gov/refseq/about/nonredundantproteins/), KOG[90], GO (http://geneontology. org/), KEGG (https://www.genome.jp/ kegg/pathway.html) and TrEMBL (http://www.bioinfo.pte.hu/ more/TrEMBL.htm) using BLAST (https://blast.ncbi.nlm.nih.gov/Blast.cgi). A total of 56,727-60,923 (92.92-97.99% of the total) protein-coding genes were annotated (**Supplementary Table 12**).

Repetitive sequences in the five species were identified using the programs LTR_FINDER[53] and RepeatScout[54]. The identified repetitive sequences were then classified using PASTEClassifier[91] and annotated using RepeatMasker (http://repeatmasker.org/). A total of 2,936,990,453- 4,145,555,209 bp Class I (67.56-71.47% of the total genome) and 515,361,078- 1,025,079,256 bp Class II (12.54-19.21% of the total genome) repetitive sequences were characterized in the five species (**Supplementary Table 13**). In the Class I, the *Copia* and *Gypsy* families consist of 16.64-22.80% and 43.72-49.58% of the assembled genome, respectively. Likewise, the Class II CACTA represent 11.06-19.21% of the assembled genome in the five species.

***Molecular phylogeny and divergence time***

Phylogenetic inferences were performed based on single copy orthologous genes (SCOGs), reduced representative genomic regions (RRGRs) and whole genome single nucleotide polymorphisms (SNPs). The SCOGs were characterized from seven diploid *Triticum*/*Aegilops* species, seven polyploid wheat subgenomes and three outgroup species, including *Triticum urartu* (abbreviated as Tura)[23], *Aegilops tauschii* (Atau)[24], *Ae. speltoides* (Aspe), *Ae. bicornis* (Abic), *Ae. longissima* (Alon), *Ae. searsii* (Asea), *Ae. sharonensis* (Asha), AA (Emaa) and BB (Emab) subgenomes of *T. turgidum*, ssp. *dicoccoides*[22], AA (Sveb) and BB (Emab) subgenomes of *T. turgidum*, ssp. *durum*[26], AA (Taea), BB (Taeb) and DD (Taed) subgenomes of *T. aestivum*[25], *Brachypodium distachyon* (Bdis)[92], *Hordeum vulgare* (Hvul)[93], *Elymus elongatum* (Ele)[94]. Homologous proteins of these diploid species and polyploid wheat subgenomes were identified using BLASTP (https://blast.ncbi.nlm.nih.gov/Blast.cgi) with E-value < $10^{e-5}$. The program OrthoFinder[73] was performed to cluster the homologous as gene families. In total, 71,986 gene families were identified from these species, of which, 38,044 contain no more than one (none or one) gene copy. We then selected 3,184 gene families (singleton) that include one copy in all these species to performed phylogenetic inference. Coding domain sequences of the singleton genes were aligned based on their amino acid sequences using Guidance [95]. A total of 2,569 gene families that generated alignments with MEAN_RES_PAIR_SCORE larger than 0.8 were retained. Then, the program Gblocks[96] was employed to further filter the low quality alignments and concatenated the high quality alignments as a single matrix. Phylogenetic tree was reconstructed using a maximum likelihood method implemented in RAxML (v.8.2.12)[57] of a GTR-GAMMA substitution model with 1,000 bootstrap replicates.

Phylogenetic relationships of the *Triticum*/*Aegilops* species were also inferred based on RRGRs using the following procedures: (1) 20-kb genic and up/down-stream regions were retrieved from the diploid species and polyploid wheat subgenomes; (2) all 20-kb genomic regions were aligned to bread wheat B-subgenome using the nucmer of MUMmer v3.9[97] with the parameters --mum -c 90 -l 40; (3) the program *delta-filter* was employed to extract one-to-one query-reference alignments with parameters -1; (4) the *show-coords* was used to obtain chromosome coordinates of each alignment; (5) the Intersect module of Bedtools (https://bedtools. readthedocs.io/en/latest/) was performed to search the intersections of alignment blocks of all query genome/subgenome regions to constructed the originally shared reduced genome on reference genome; (6) the reduced reference genome sequences were aligned against to their own reference genomes using Minimap2[74] with parameter "–x asm20". In total, we obtained ~30 Mb reduce genome sequences, which contains 2,223 reduced genomic regions. In addition, we also reconstructed phylogenies of the *Triticum*/*Aegilops* species based on whole genome resequencing data. The clean short reads of seven diploid *Triticum*/*Aegilops* species were mapped onto bread wheat B-subgenome using BWA (http://bio-bwa.sourceforge.net/). Genome-wide SNPs were reported using SAMtools (http://www.htslib.org/). Phylogenetic tree was reconstructed using a maximum likelihood method implemented in RAxML (v.8.2.12)[57] of a GTR-GAMMA substitution model with 1,000 bootstrap replicates.

Divergence times among these *Triticum*/*Aegilops* species were estimated based on SCOGs, RRGRs and genome-wide SNPs using the program BEAST (v.2.6.0)[58] with the following parameters, strict molecular clock, HKY + gamma nucleotide substitution model, Yule priors and running for 100 million MCMC generations with parameters sampled every 10000 generations. The previously estimated divergence time between *B. distachyon* and *Triticum*/*Aegilops* (mean 44.4 ± stdev 3.53 Ma)[4] was used as date calibration. TreeAnnotator (v.2.6.0) was used to summarize the output. The resulting phylogenies were visualized with ggtree[59].

#### *Genome-wide genetic similarity*

Genome-wide similarity was estimated based on the above generated RRGRs. Pair-wise nucleotide diversity between any two of these diploid species and polyploid wheat subgenomes were calculated using the DendroPy module[61] in python. Maximum likelihood tree was reconstructed using RAxML[57] with GTR-GAMMA model. The R package Ape v5.1[98] was used to manipulate phylogenetic trees with the following parameters: chronos function was used to make ultrametric trees from input trees; conphenetic.phylo function computed the pairwise branch lengths between any pairs of phylogenetic tips; keep.tip function extracted needed tips and generate local tree topologies; tree topology was visualized with plot.phylo function. The Treedist function of R package phangorn was employed to calculate the difference between phylogeny trees to classify tree topologies of each reduced representative genome region. Tree topologies along each chromosome were visualized with RIdeogram (https://github.com/TickingClock1992/RIdeogram).

***Estimation of genetic introgression***

Reduced representative genomic regions of the nine diploid species and bread wheat B-subgenome were employed to estimate introgression events. Given a quartet of three species and a constant outgroup (*Elymus elongatum*) with the relationship (((P1, P2), P3), O) corresponding to *Triticum*/*Aegilops* phylogeny topology, we performed different statistics, such as *D*, *fd*, hybrid index ($\gamma$) and $\chi^2$ goodness of fit test, to systematically infer the introgression/hybridization events among the selected *Triticum*/*Aegilops* species. The statistics *D* and *fd* were calculated with genomics_general tools[62] (https://github.com/simonhmartin/genomics_general). We used a significance threshold of 0.001 for the *Z*-test. The program HyDe[63] was used to calculate the hybrid index ($\gamma$), which quantifies the proportion of P3 contributed to P2. In addition, $\chi 2$ goodness of fit test is also applied to identify genetic introgressions based on the counts of three different sliding-window trees, with the (((P1, P2), P3) representing the topology of species tree and ((P2, P3), P1) and ((P1, P3), P2) representing discordant trees[64]. Ratios between two discordant tree topologies that are significantly deviated from 1 ($p$-value < 0.001) indicate the presence of introgression event.

***Evolutionary history of the gene families***

To obtain the core protein-coding gene set of the *Triticum*/*Aegilops* species, we clustered the homologous into gene families using OrthoFinder[73]. All the three polyploid wheats (wild emmer, durum and bread wheat) and their A- and D-subgenome donors, *T. urartu* and *Ae. tauschii*, were combined as an *in-silico* species called "ABD". Then, the *in-silico* species and five Sitopsis species were used to construct presence/absence matrix according to the member count in each family (presence marked as "1" whereas absence marked as "0"). In total, 30,469 gene families corresponding to three categories were reconstructed, including: (1) core gene family, present in all *Sitopsis* species; (2) dispensable gene family, present in either at least two (but not all) *Sitopsis* species or the ABD with at least one (not all) *Sitopsis* species; (3) *Sitopsis*-specific gene family, present only in one *Sitopsis* species. Intersect analyses of the three gene categories were visualized by R package Upset (https://github.com/hms-dbmi/UpSetR/).

We also evaluated the gene family expansion and contraction histories of the *Triticum*/*Aegilops* species using a previously published phylogenomic approach[25] with the same protein-coding gene set. Log-transformed gene family size of these *Triticum*/*Aegilops* species were compared using the phylANOVA function of phytools package (https://github. com/liamrevell/phytools) in R. The *p*-value of ANOVA statistics was corrected using FDR method. Only these gene families that possess FDR < 0.1 were applied to infer the expansion and contraction history.

***Annotation of NBS-LRR and agronomic/domestication trait-related genes***

Annotation of NBS-LRR genes of the *Triticum*/*Aegilops* species were performed using NLR-Annotator pipeline[77]. The program PfamScan (https://www.ebi.ac.uk/Tools/pfa/pfamscan/) was used to search the candidate NBS-LRR genes against Pfam database (https://pfam.xfam.org/) to validate the integrity of NB_ARC domain (PF00931). All identified NBS-LRR genes were classified according to the position of intron located inside or outside of NB_ARC domain. In addition, we also employed the available coding sequences of agronomic/domestication trait-related gene (*i.e*., *Q/q* gene) as query to search against to *Triticum*/*Aegilops* reference genomes using BLASTN with e-value < $10^{e-5}$ and hits > 300bp in length. All candidate hits were manually checked and compared with annotated protein sequences. For BLAST hits that missed in annotated protein sequences, DNA segments together with flanking regions of 200 bp were extracted and annotated using both ORFfinder (https://www.ncbi.nlm. nih.gov/orffinder/) and Augustus (http://bioinf.uni-greifswald.de/augus tus/submission.php). Newly annotated genes were BLAST against original gene set for validation.

**Supplementary Figures:**

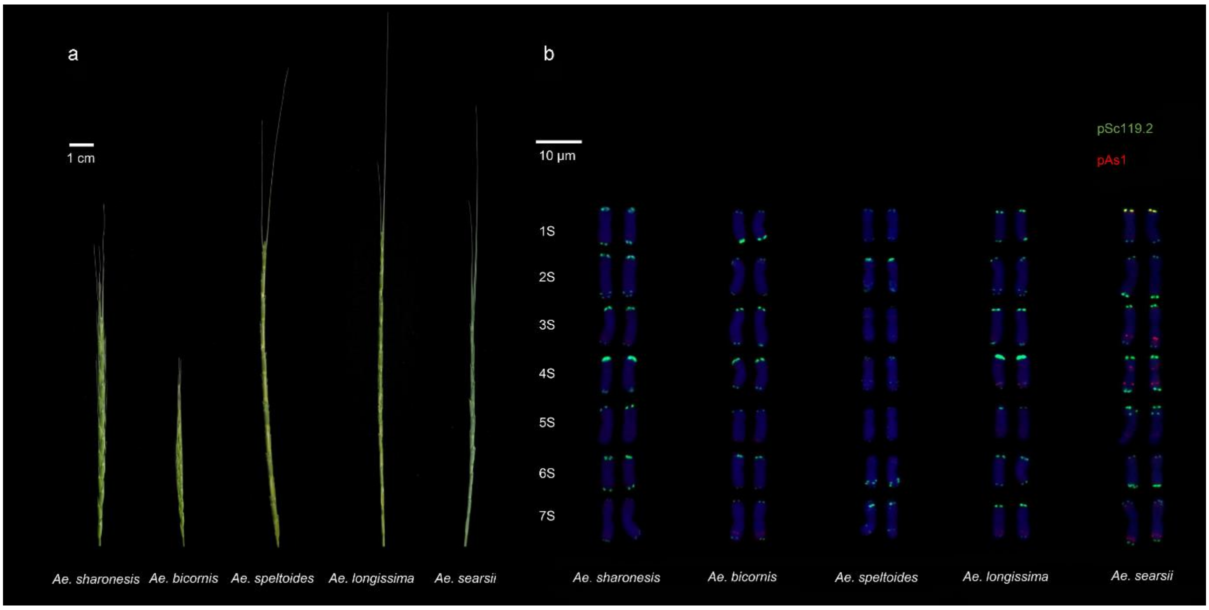


**Supplementary Fig.1**. Spike morphology (**a**) and fluorescence in situ hybridization (FISH) (**b**) of the five *Sitopsis* species. The same accessions of the five species were used to conduct *de novo* assembly.

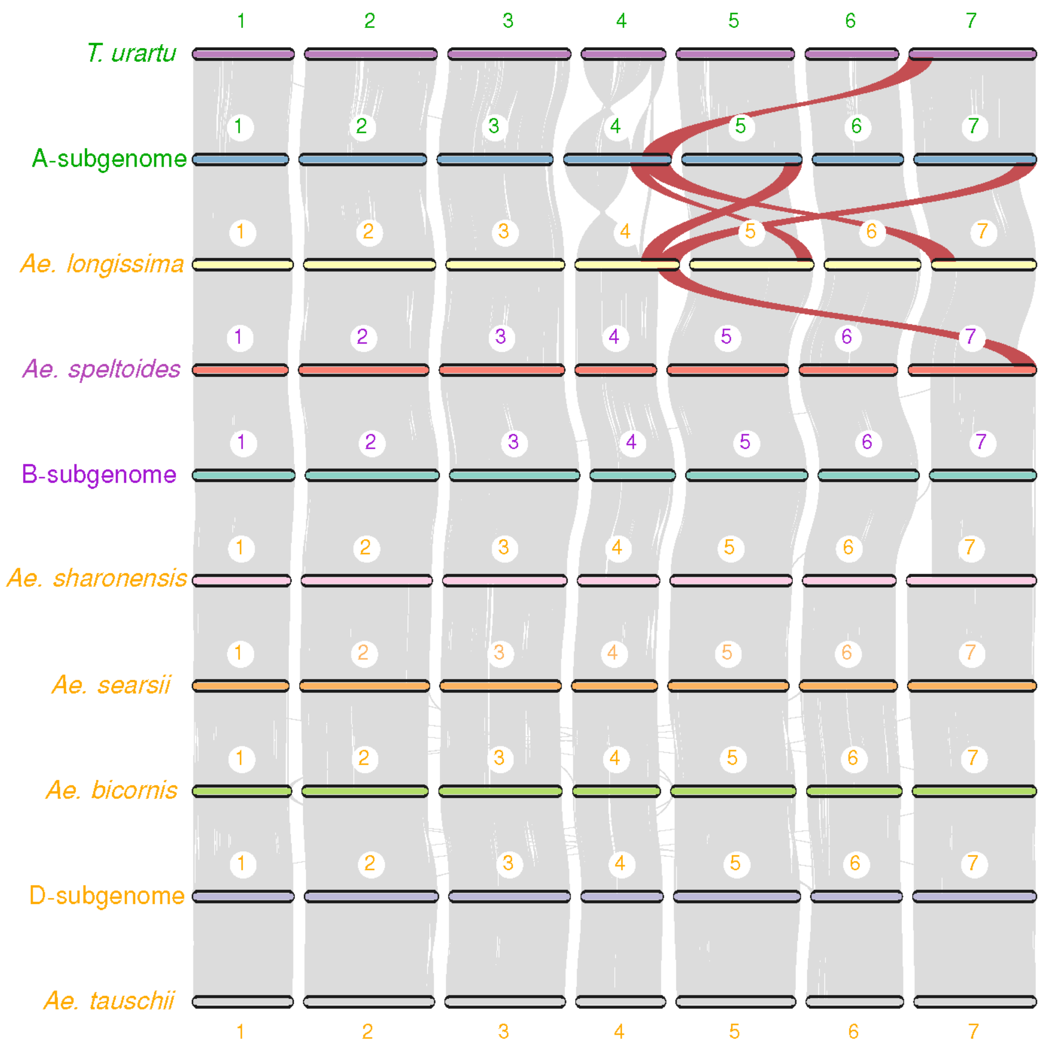


**Supplementary Fig. 2**. Genome collinearity analysis of the *Triticum*/*Aegilops* species complex. The two previously identified large genomic translocations (4A/5A/7B in bread wheat and $4^{l}/7^{l}$ in *Ae. longissima*) are highlighted by red color.

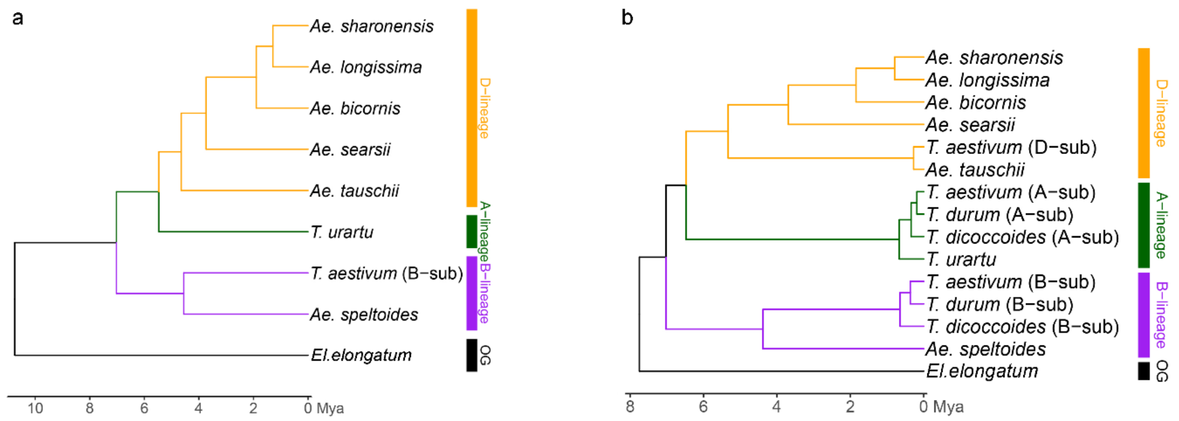


**Supplementary Fig. 3**. Maximum likelihood tree and divergence time of the *Triticum*/*Aegilops* species based on whole genome SNPs (a) and reduced representative genomic regions (b).

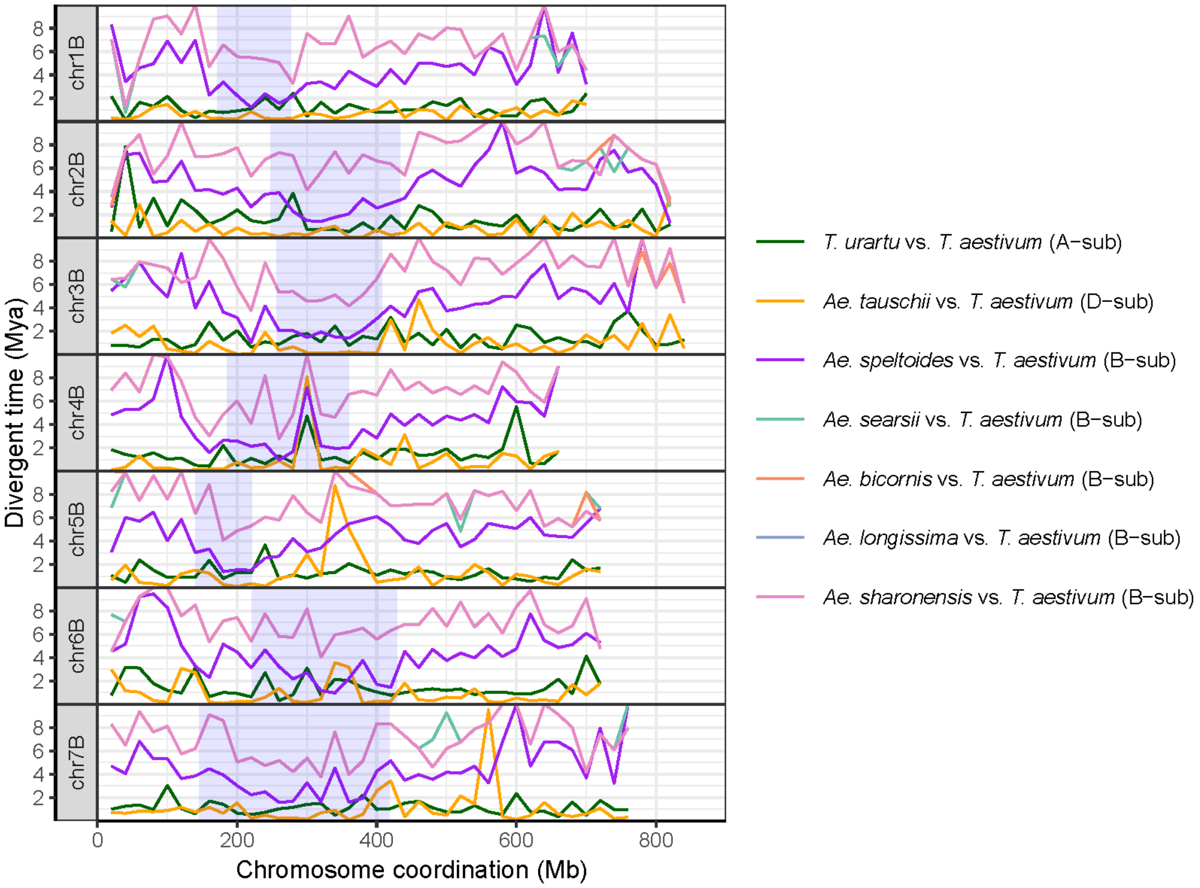


**Supplementary Fig. 4**. Divergence time between the seven diploid *Triticum*/*Aegilops* species and polyploid wheat B-subgenome along the seven chromosomes based on single copy ortholog genes.

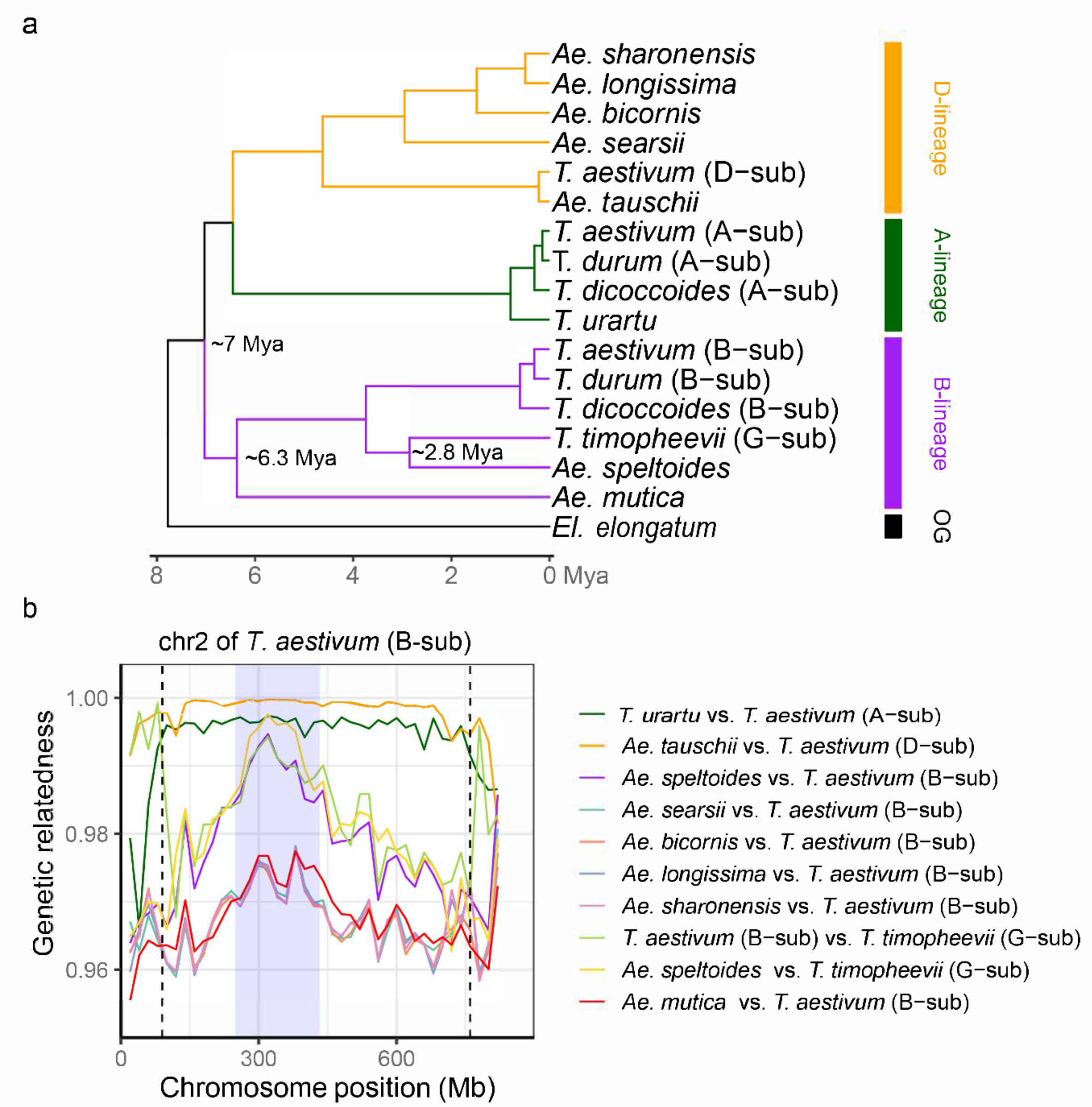


**Supplementary Fig. 5**. Maximum likelihood tree (a) and pair-wise comparisons of the genetic relatedness (b) between the seven diploid *Triticum*/*Aegilops* species and bread wheat B-subgenome and Timopheevii G-subgenome based on the reduced representative genomic regions of chromosome 2.

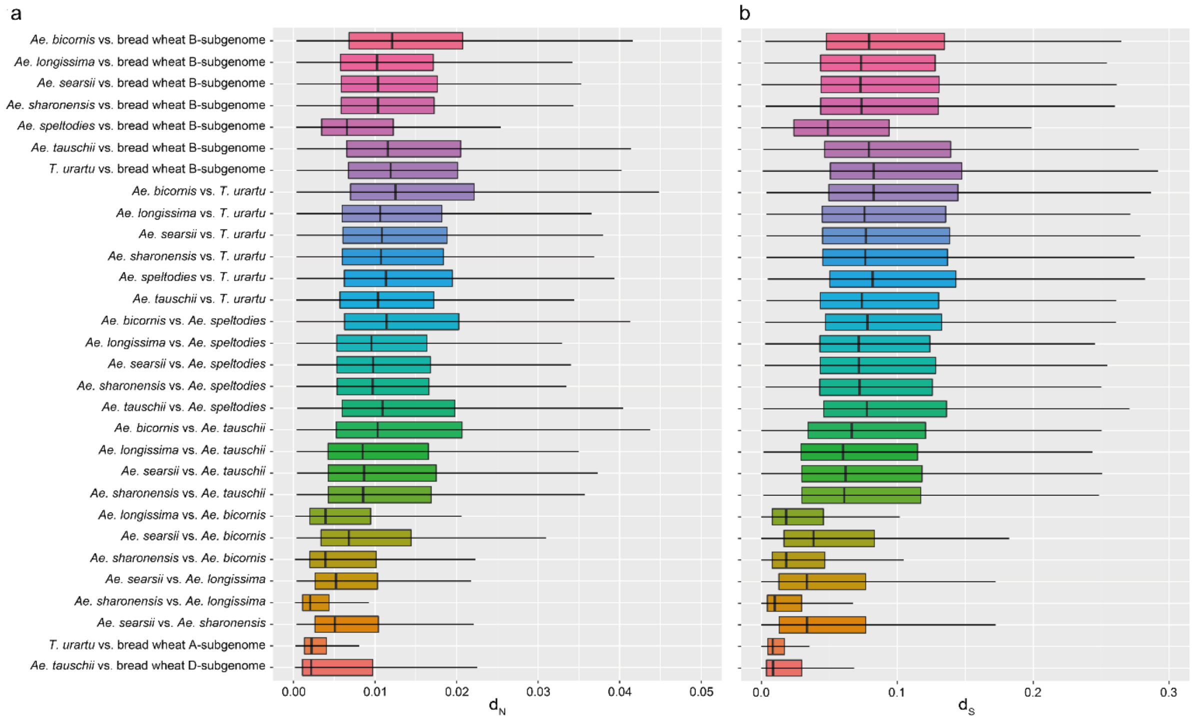


**Supplementary Fig. 6**. Pair-wise nonsynonymous ($d_N$) and synonymous ($d_S$) substitution rates of the diploid *Triticum*/*Aegilops* species and polyploid wheat subgenomes based on single copy orthologous genes.

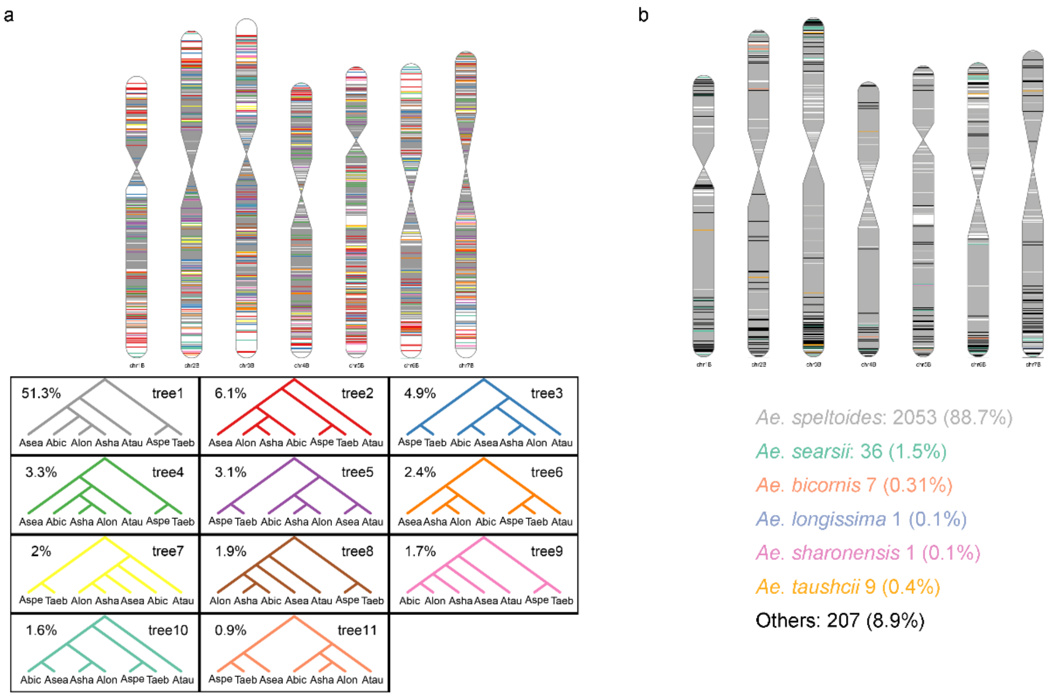


Taeb: *T. aestivum* (B-sub); Aspe: *Ae. speltoides*; Asea: *Ae. searsii*; Abic: *Ae. bicornis*; Alon: *Ae. longissima*; Asha: *Ae. sharonensis*; Atau: *Ae.tauschii*

**Supplementary Fig. 7**. Distribution of the tree topologies of the *Triticum*/*Aegilops* species based on 2,314 reduced representative genomic regions. (a) Only the top 11 tree topologies were shown. Colors on the seven bread wheat B-subgenome chromosomes are the same as the tree topologies. (b) Distribution and percentages of the incongruent gene tree between the five *Sitopsis* species and their closely relatives. Colors on the seven bread wheat B-subgenome chromosomes are the same as the species names.

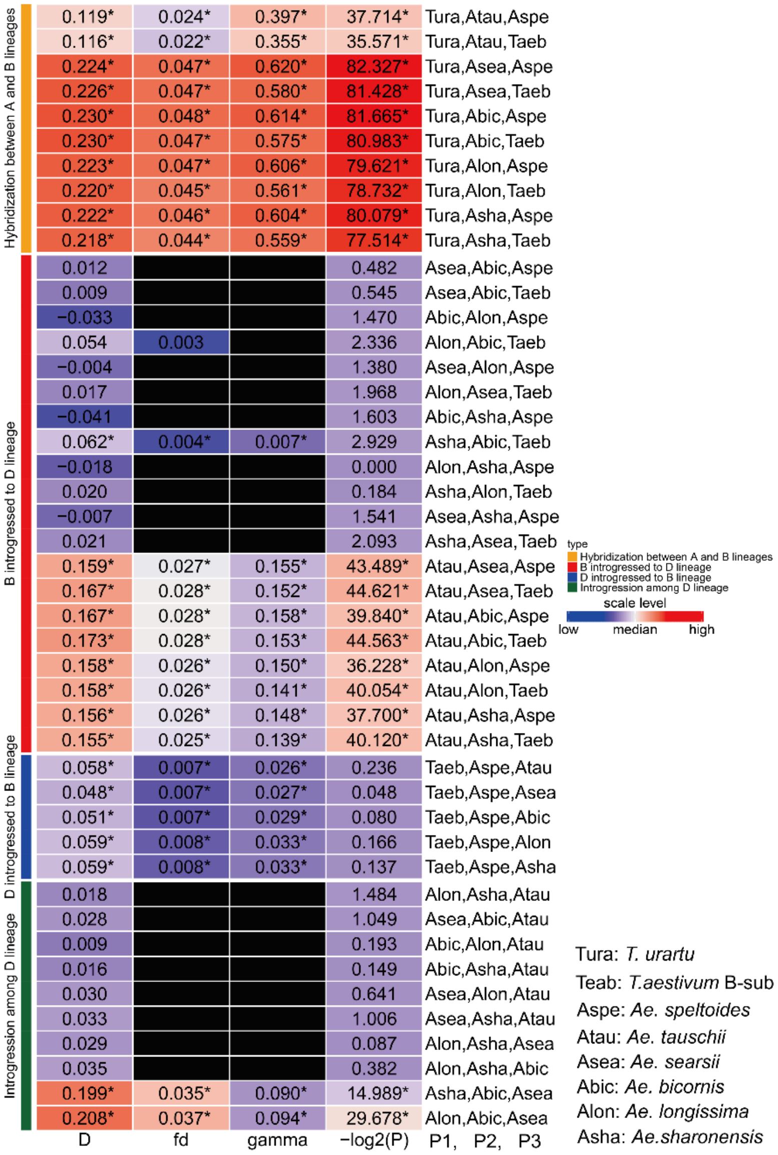


**Supplementary Fig. 8**. Estimates of the *fd* and *D* value among the *Triticum*/*Aegilops* species based on the representative genomic regions.

a

| | | *T. urartu* (A) | wheat B-subgenome (B) | *Ae. speltoides* (Aspe) | Ancestor of B and Aspe | *Ae. tauschii* |
|---|---|---|---|---|---|---|
| Specific sites shared with n *Sitopsis* species | 4 | 44,580 | 20,749 | 20,665 | 61,104 | 232,995 |
| | 3 | 25,165 | 18,271 | 17,940 | 25,451 | 55,624 |
| | 2 | 13,565 | 14,992 | 12,957 | 14,505 | 19,513 |
| | 1 | 45,960 | 47,908 | 36,694 | 40,150 | 62,771 |
| | Total | 129,270 | 101,920 | 88,256 | 141,210 | 370,903 |
| | Pro. | 1.31% | 1.04% | 0.90% | 1.44% | 3.77% |
| Specific sites shared with *Ae. tauschii* | Num | 87,963 | 25,380 | 25,788 | 29,235 | N |
| | Pro. | 0.89% | 0.26% | 0.26% | 0.30% | N |

b

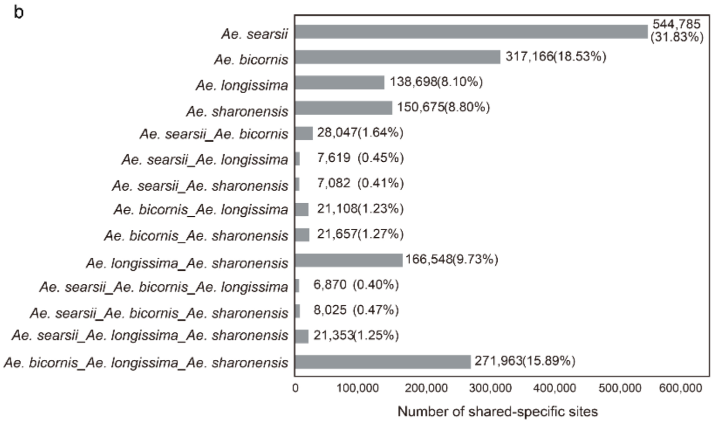


**Supplementary Fig. 9**. Percentages of the putative introgressed sites (a) and shared species-specific SNPs (b) among the *Triticum*/*Aegilops* species.

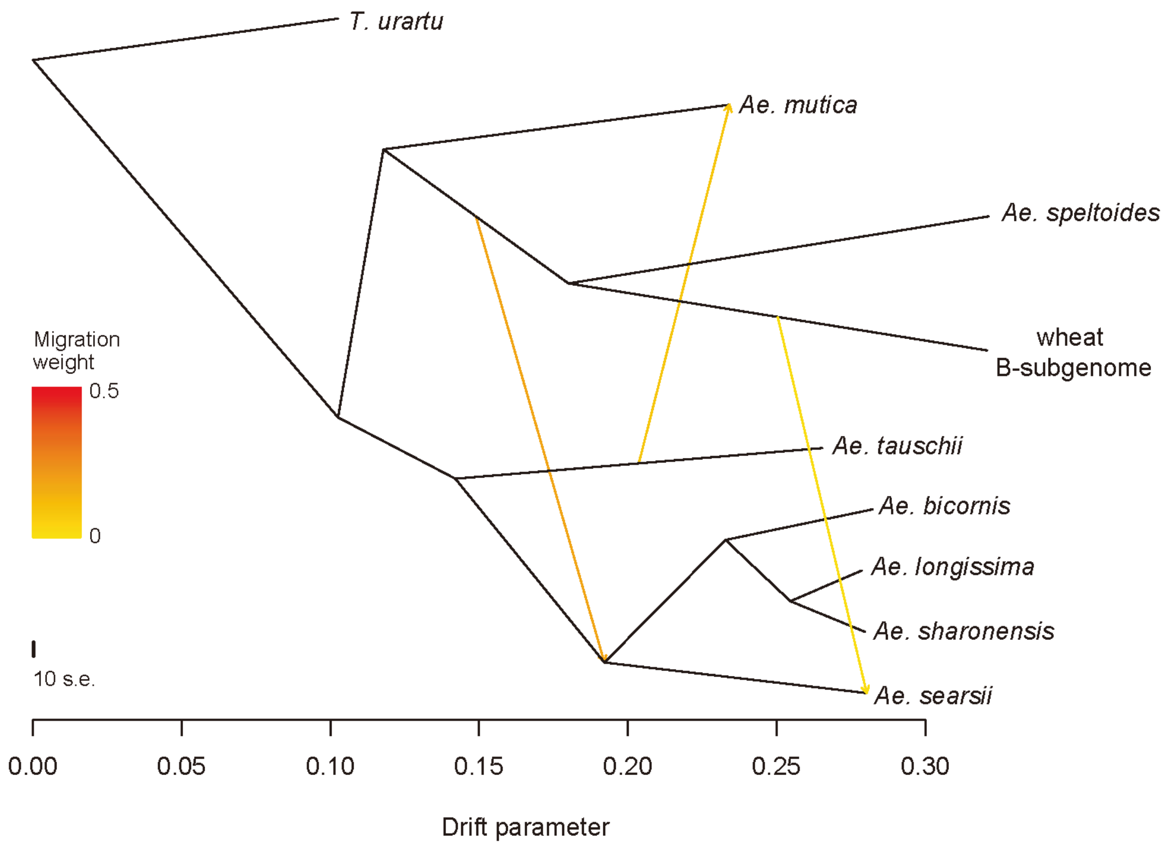


**Supplementary Fig. 10**. Allele frequency-based migration inference of the seven diploid *Triticum*/*Aegilops* species and bread wheat B-subgenome using Treemix based on whole genome SNPs. Arrows with color from orange to red indicate the high and low possibility migration events. Only the top three migration events are shown. Bar on the bottom represents the branch length of maximum likelihood tree.

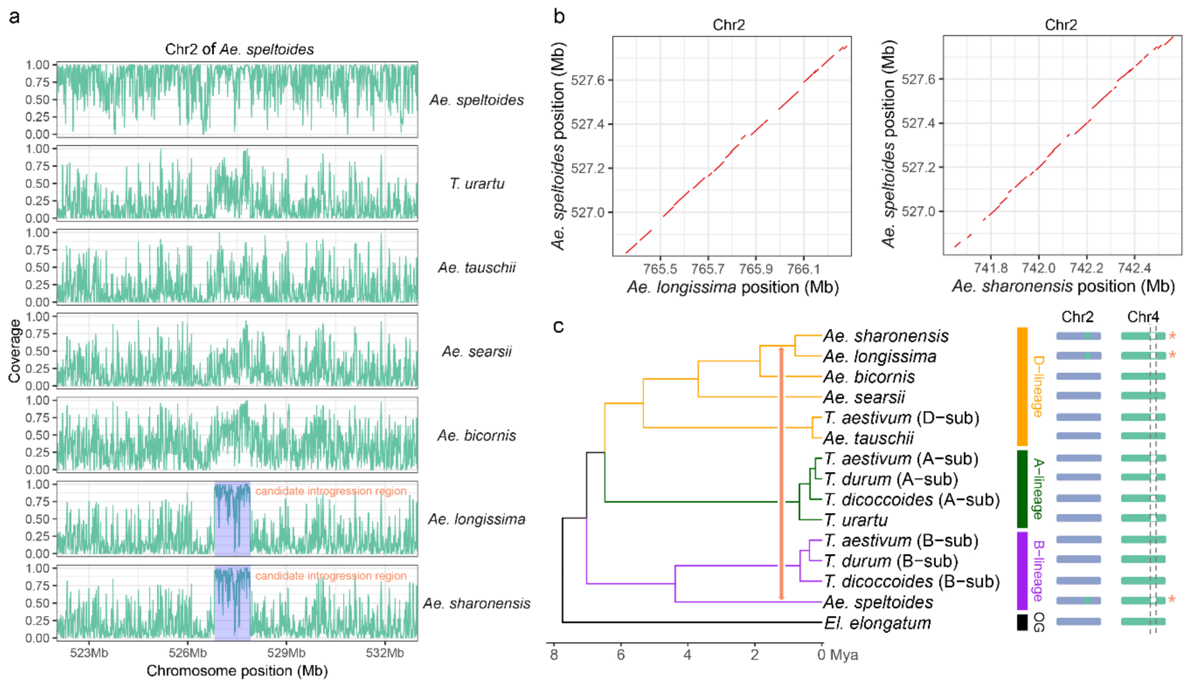


**Supplementary Fig. 11**. Putative introgressed genomic regions between the *Ae. speltoides* and the D-lineage *Sitopsis* species based on the inter-specific sequence similarity through whole-genome alignment and mapping coverage of Illumina short reads. (a) Putative introgressed genomic region identified between both the *Ae. longissima* and *Ae. sharonensis* and *Ae. speltoides*. This genomic region was identified through genome-wide inter-specific sequence alignment. X indicates the coordinate on *Ae. speltoides* chromosome 2. Y axis represents the total reads of the seven diploid species mapped onto the *Ae. speltoides* reference genome. Only the *Ae. longissima* and *Ae. sharonensis* show high read coverage at this genomic region. (b) Dotplots indicate the high collinear genomic region on chromosome 2 between *Ae. speltoides* and each of the *Ae. longissima* and *Ae. sharonensis*. (c) The genomic region on chromosome 2 (green color) was translocated from chromosome 4 (white color). The same genomic region shared in the *Triticum*/*Aegilops* species is most likely through inter-specific genetic introgression.

a

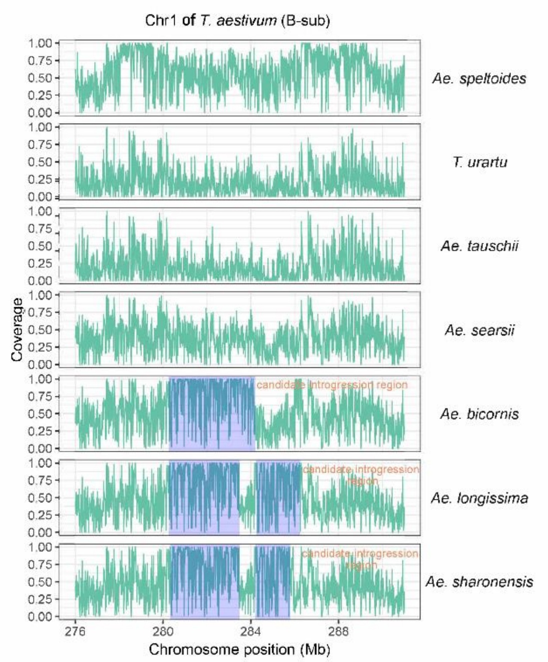


b

Introgression events among Sitopsis species
(0. not occurred; 1. occurred)

| Event id | *Ae. searsii* | *Ae. bicornis* | *Ae. longissima* | *Ae. sharonensis* |
|---|---|---|---|---|
| 1 | 0 | 1 | 1 | 1 |
| 2 | 1 | 0 | 0 | 0 |
| 3 | 1 | 1 | 0 | 0 |
| 4 | 1 | 0 | 0 | 0 |
| 5 | 1 | 1 | 1 | 1 |
| 6 | 1 | 1 | 1 | 1 |
| 7 | 1 | 1 | 0 | 0 |
| 8 | 1 | 1 | 1 | 1 |
| 9 | 1 | 0 | 0 | 0 |

**Supplementary Fig. 12**. Putative introgressed genomic regions between bread wheat B-subgenome and the four D-lineage *Sitopsis* species based on the inter-specific sequence similarity through whole-genome alignment and mapping coverage of Illumina short reads. On the left, X indicates the coordinate on B-subgenome chromosome 1. Y axis represents the total reads of the seven diploid species mapped onto the B-subgenome reference genome. On the right, the red number 1 (highlighted by black color) represents the putative introgression event between bread wheat B-subgenome and the four D-lineage *Sitopsis* species.

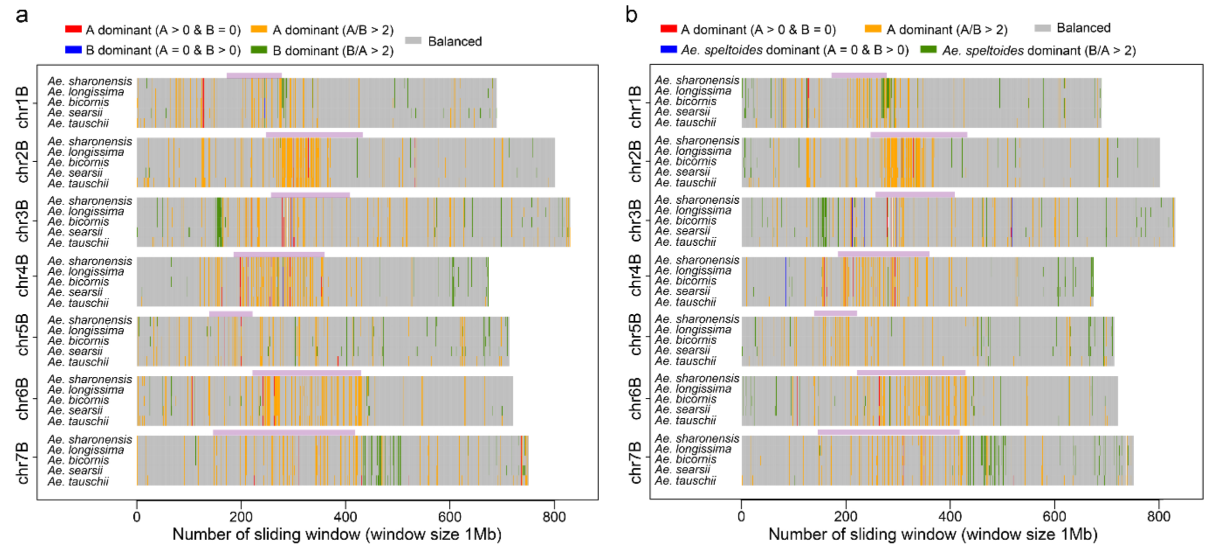


**Supplementary Fig. 13**. Distribution patterns of the A- and B-lineage specific SNPs in the five D-lineage species. The A-dominant genomic regions are defined as the total number of A-lineage specific SNPs are >2 fold higher than B-lineage specific SNPs (or no B-lineage specific SNPs). The reverse pattern is defined as B-dominant genomic region. The remaining genomic regions that are neither A- nor B-lineage dominant are defined as balanced genomic regions.

**a**

| Species name | *gypsy* | *copia* | *CACTA* | Class I other | Class II other | Unclassified | All repeats | Genome size |
|---|---|---|---|---|---|---|---|---|
| *T. aestivum* (A-sub) | 2.48 | 0.88 | 0.64 | 0.19 | 0.07 | 0.04 | 4.30 | 4.93 |
| *T. dicoccoides* (A-sub) | 2.46 | 0.88 | 0.64 | 0.19 | 0.07 | 0.04 | 4.28 | 4.90 |
| *T. durum* (A-sub) | 2.44 | 0.87 | 0.64 | 0.19 | 0.07 | 0.04 | 4.25 | 4.86 |
| *T. aestivum* (B-sub) | 2.41 | 0.86 | 0.80 | 0.25 | 0.08 | 0.07 | 4.48 | 5.18 |
| *T. dicoccoides* (B-sub) | 2.42 | 0.86 | 0.81 | 0.25 | 0.08 | 0.06 | 4.49 | 5.18 |
| *T. durum* (B-sub) | 2.39 | 0.85 | 0.79 | 0.25 | 0.08 | 0.06 | 4.42 | 5.11 |
| *Ae. speltoides* | 1.80 | 0.94 | 0.45 | 0.20 | 0.06 | 0.09 | 3.54 | 4.11 |
| *T. aestivum* (D-sub) | 1.64 | 0.66 | 0.77 | 0.20 | 0.07 | 0.04 | 3.38 | 3.95 |
| *Ae. tauschii* | 1.70 | 0.71 | 0.82 | 0.21 | 0.07 | 0.05 | 3.56 | 4.22 |
| *Ae. longissima* | 2.86 | 1.03 | 0.81 | 0.26 | 0.08 | 0.07 | 5.11 | 5.80 |
| *Ae. sharonensis* | 2.92 | 1.03 | 0.83 | 0.26 | 0.08 | 0.08 | 5.19 | 5.89 |
| *Ae. searsii* | 2.39 | 0.89 | 0.95 | 0.27 | 0.08 | 0.06 | 4.64 | 5.34 |
| *Ae. bicornis* | 2.72 | 0.98 | 0.92 | 0.29 | 0.09 | 0.08 | 5.08 | 5.78 |

**b**

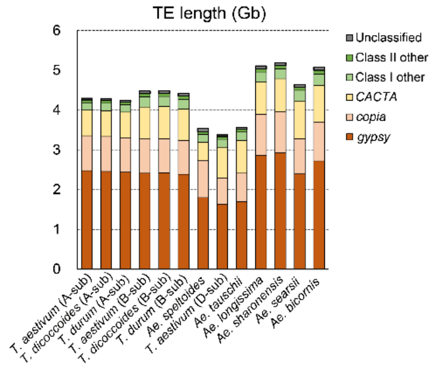


**c**

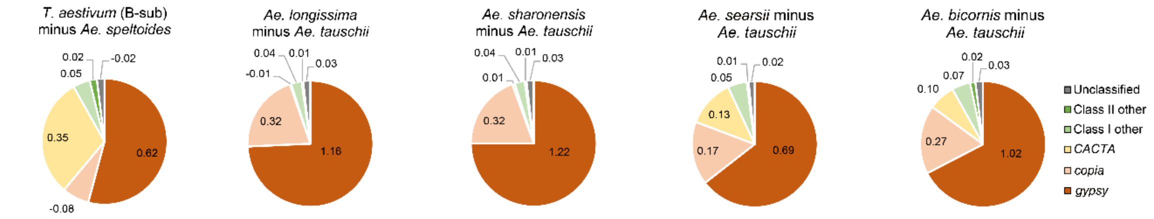


**Supplementary Fig. 14**. Genome-wide feature of the repetitive sequence in the diploid species and polyploid wheat subgenomes. (a-b) Total length of each type of the repetitive sequence. (c) Difference in total length of the repetitive sequence in the B-lineage (between *Ae. speltoides* and bread wheat B-subgenome) and D-lineage (between *Ae. tauschii* and the four D-lineage *Sitopsis* species).

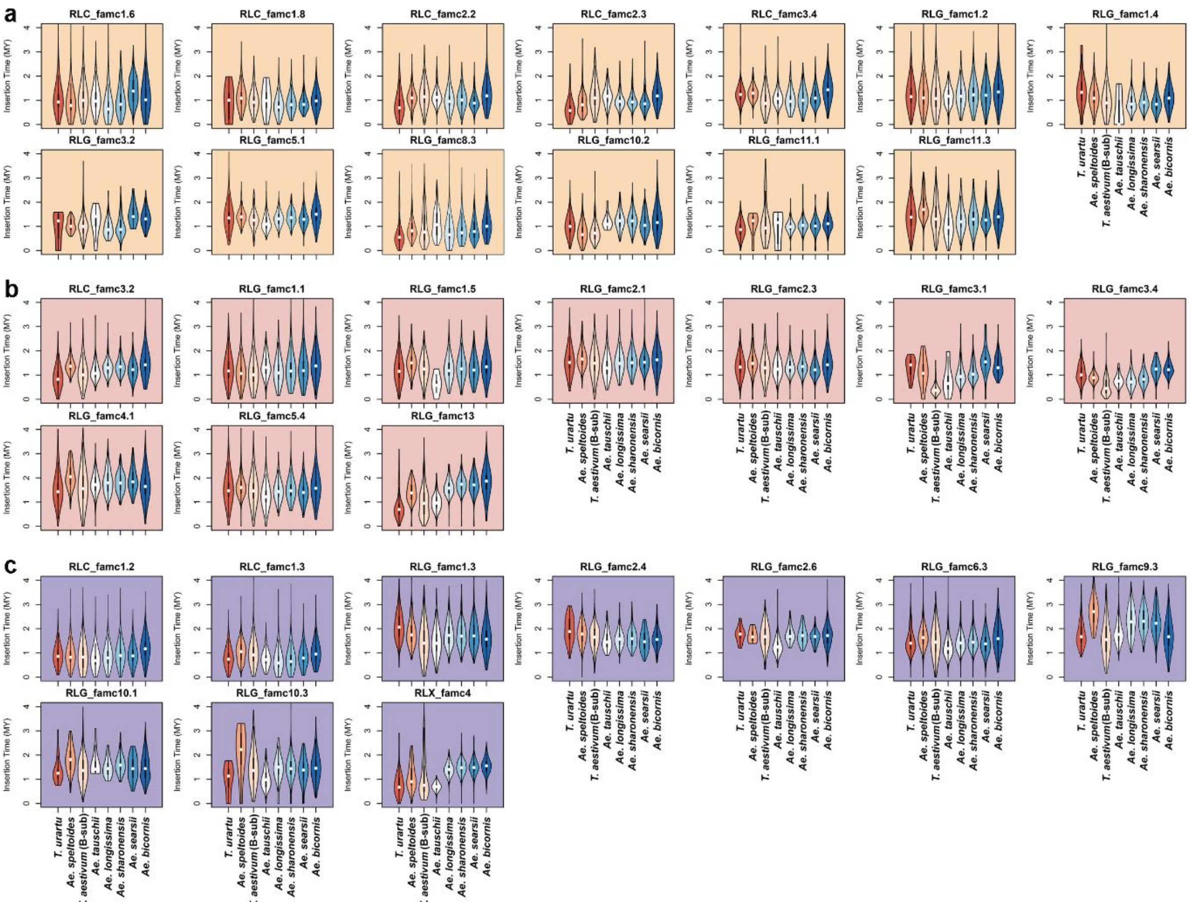


**Supplementary Fig. 15**. Insertion times of the transposon subfamilies that amplified specifically in D- (orange) and B-lineages (purple) (a and c) and shared between the two lineages (red) (b).

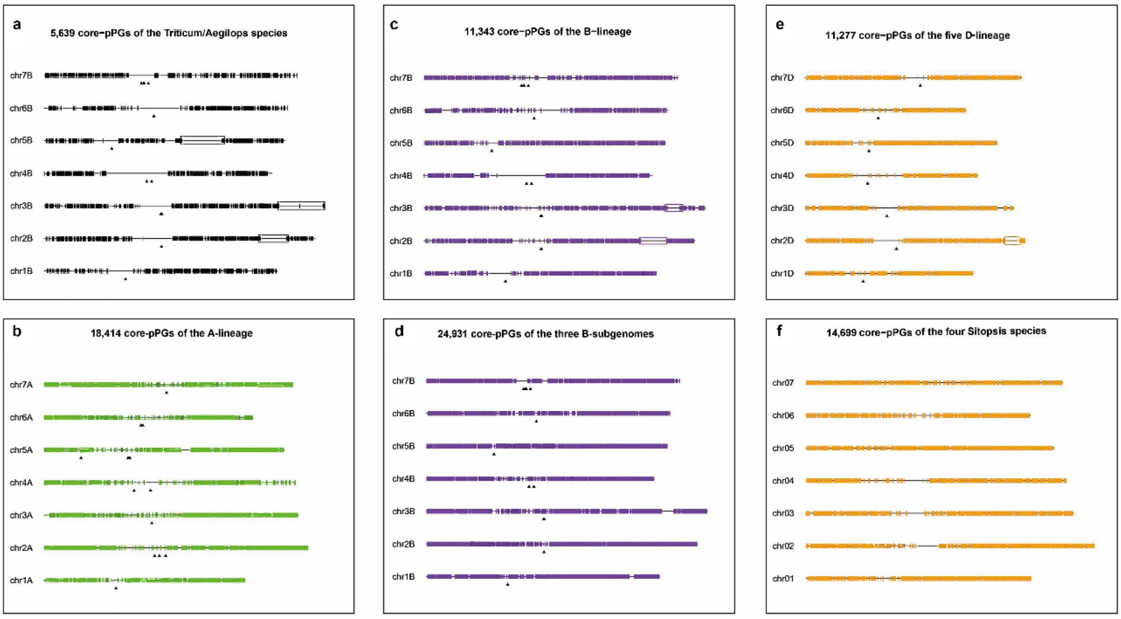


**Supplementary Fig. 16**. Distribution patterns of the core putative proto-gene (pPG) in all the *Triticum*/*Aegilops* species complex (black) (a), A-lineage (green) (b), B-lineage (purple) (c), B-subgenome (purple) (d), D-lineage (orange) (e), and D-lineage *Sitopsis* species (orange) (f). Identification of the core pPG in the D-lineage *Sitopsis* species were defined based on the *Ae. longissima* reference genome, all the other core pPG were defined based on the three subgenomes of bread wheat. Non-conserved genomic regions in the B-, D- and all *Triticum*/*Aegilops* species are shown in purple, orange and black boxes.

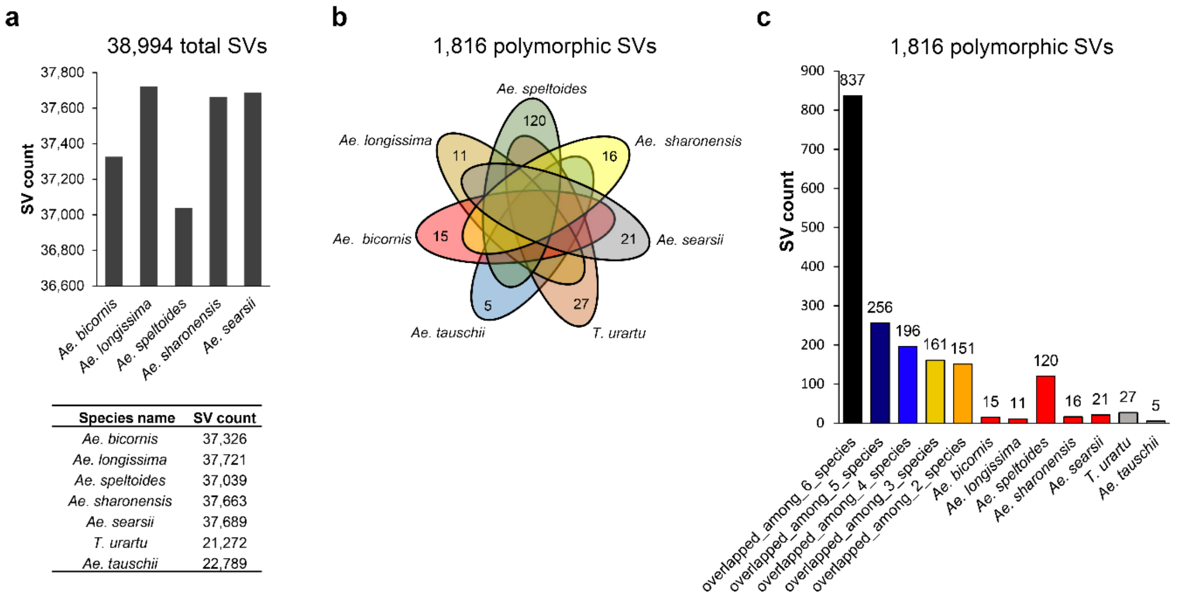


**Supplementary Fig. 17**. Pan-genomic analyses of the genome structural variations (SVs) of the seven diploid *Triticum*/*Aegilops* species. (a) Total numbers of SVs identified in the seven diploid species. (b-c) Intersection analysis of the 18,16 polymorphic SVs among the seven diploid species.

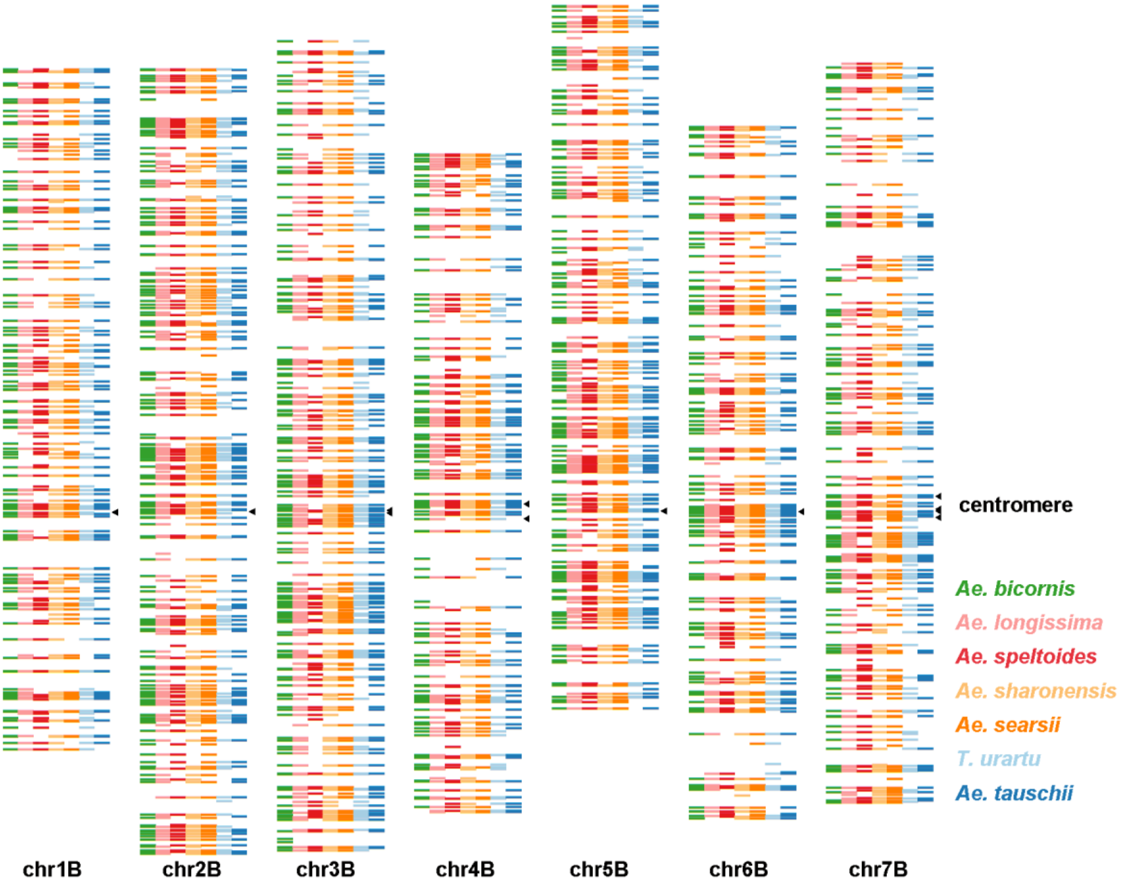


**Supplementary Fig. 18**. Genome-wide distribution of the 1,816 shared polymorphic SVs in the seven diploid species on the seven chromosomes.

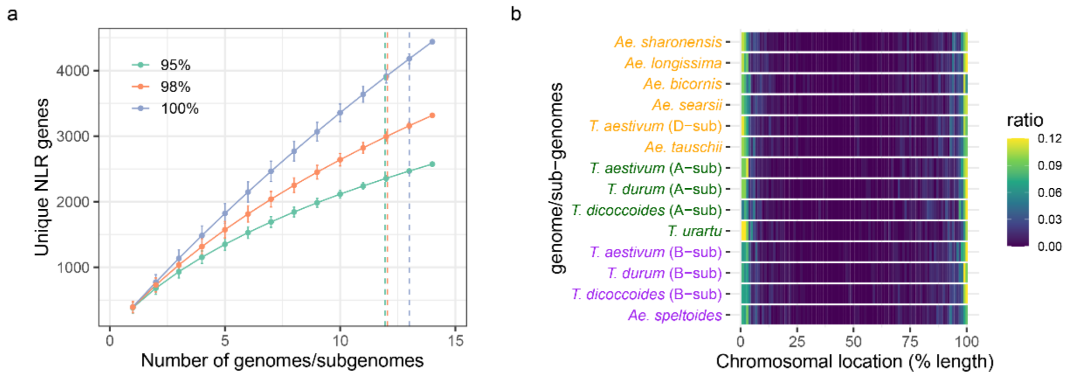


**Supplementary Fig. 19**. Identification of nucleotide-binding and leucine-rich repeat (NBS-LRR) gene in the *Triticum*/*Aegilops* species. (a) Numbers of unique NBS-LRR genes identified in the *Triticum*/*Aegilops* species based on 95%, 98% and 100% genetic identity. X and Y-axes indicate the numbers of genomes and NBS-LRR genes. (b) Genome-wide distribution of the of the NBS-LRR genes in the *Triticum*/*Aegilops* species.

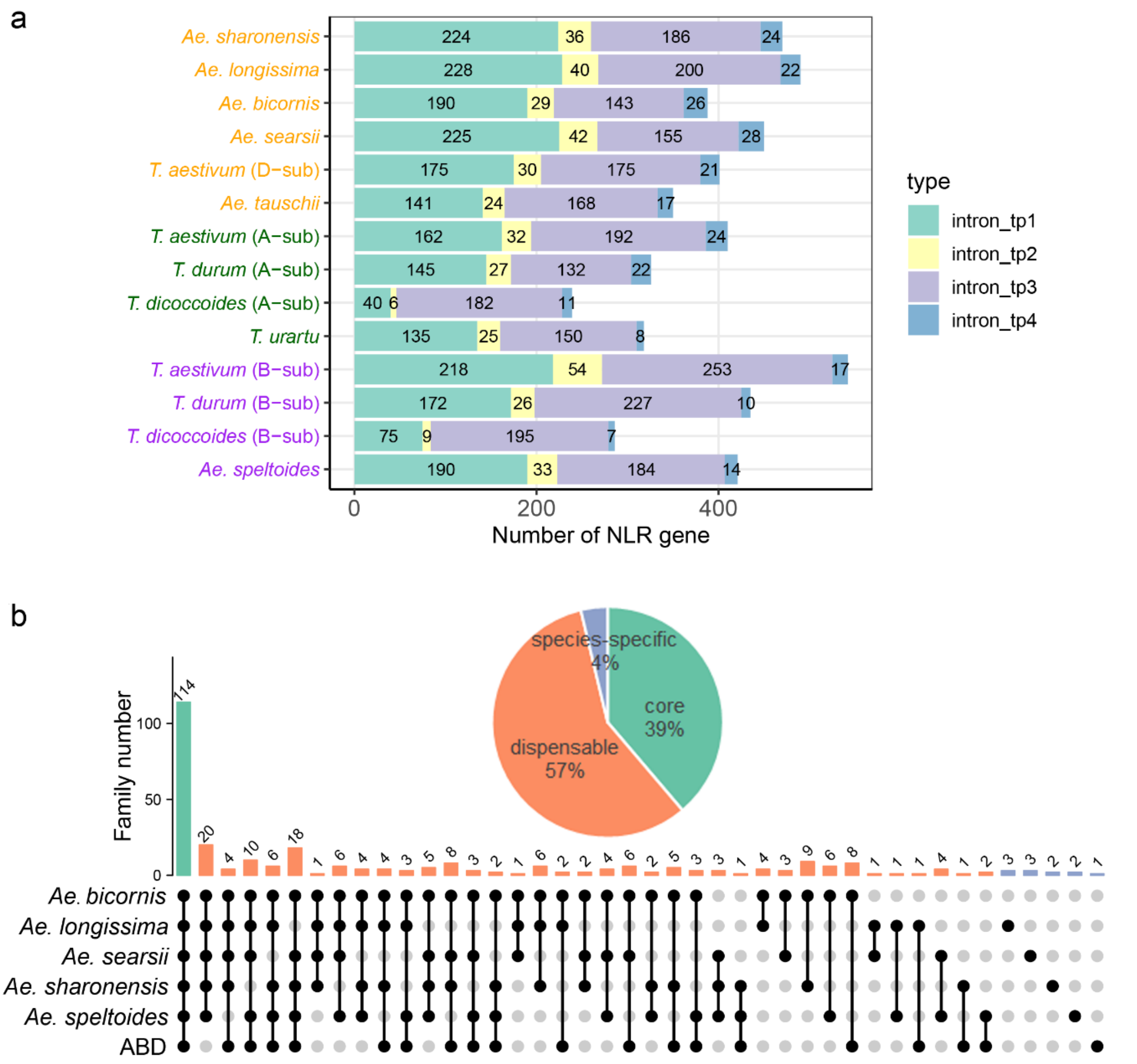


**Supplementary Fig. 20**. (a) Total NBS-LRR genes identified in the diploid species and polyploid wheat subgenomes. Tuiquoise, yellow, orchid and blue colors indicate the different types of NBS-LRR genes. (b) Intersection analysis of the NBS-LRR genes in the *Triticum*/*Aegilops* species.

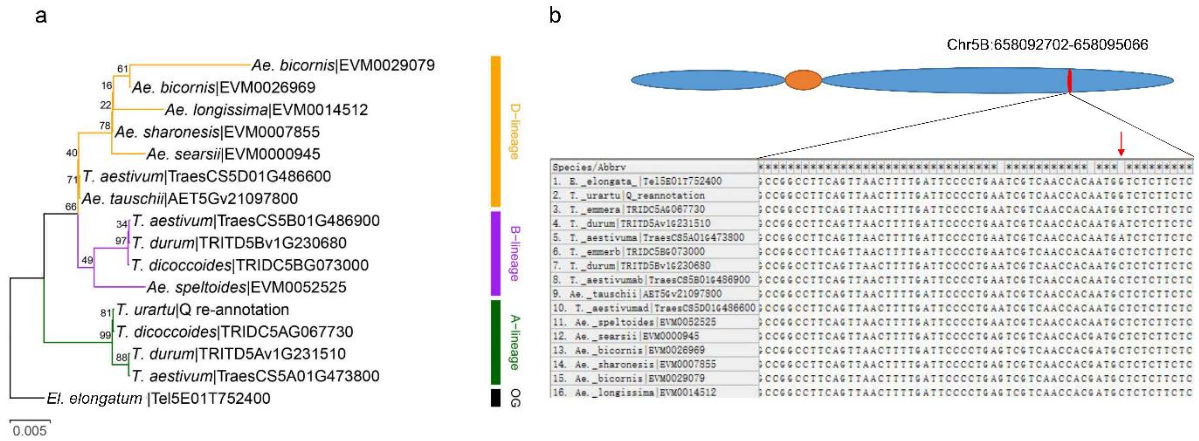


**Supplementary Fig. 21**. Phylogenetic tree (a) and domestication allele (b) of the *Q*/*q* gene in in the *Triticum*/*Aegilops* species. The red arrow indicates the key mutation of the *Q* ($I_{329}$) and *q* ($L_{329}$ and $V_{329}$) alleles.

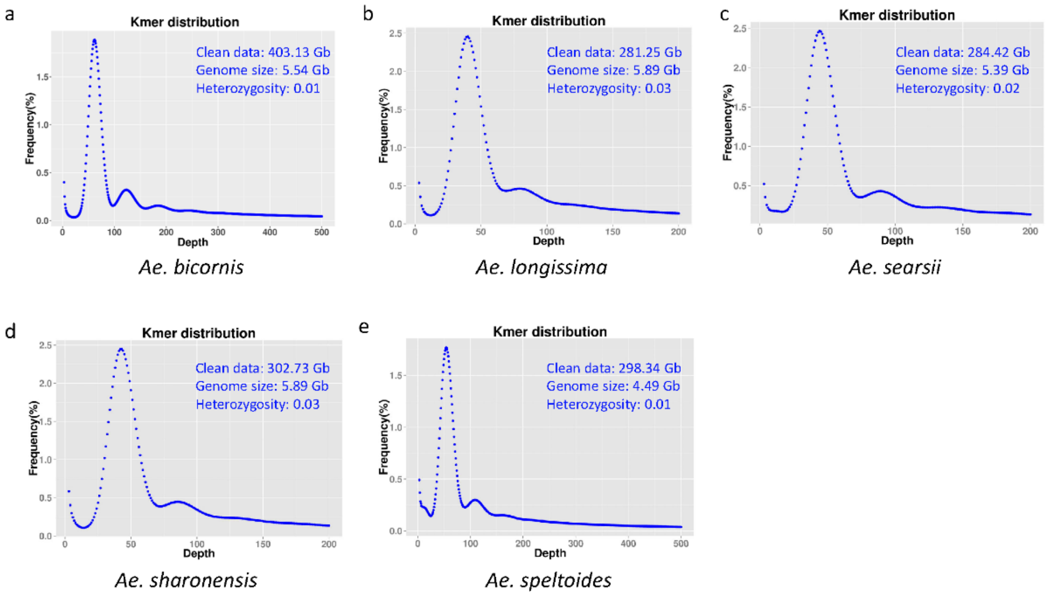


**Supplementary Fig. 22**. Genome survey of the five *Sitopsis* species based on Illumina short reads.

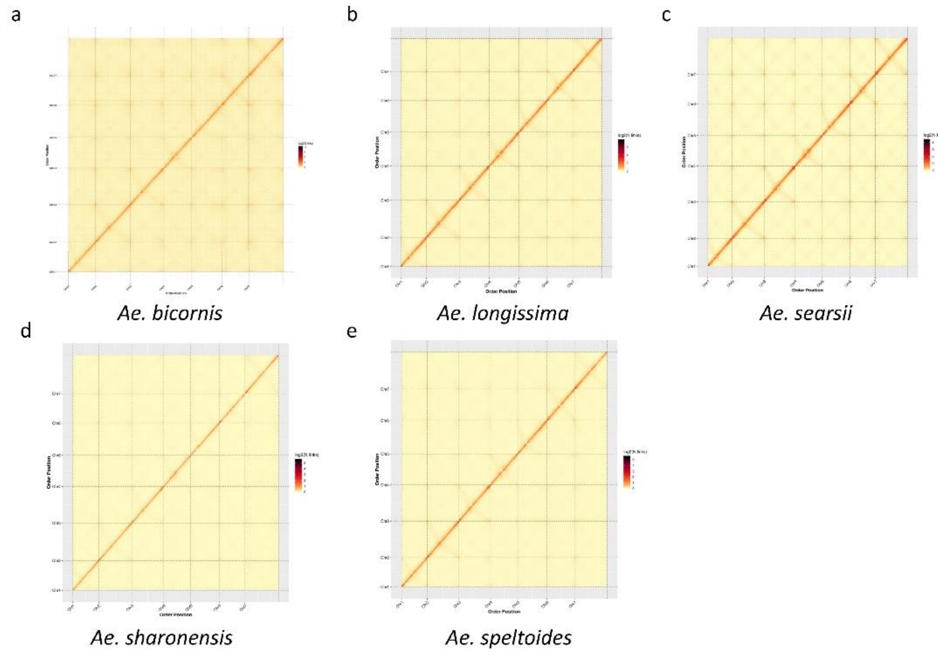


**Supplementary Fig. 23**. Genome-wide analysis of chromatin interactions at 500-kb resolution of the five *Sitopsis* species.
